# Genetically engineered rapamycin responsive K2P channels

**DOI:** 10.64898/2026.05.21.726927

**Authors:** Leila Khajoueinejad, Elena B. Riel, Aben Ovung, Aboubacar Wague, Caroline S. Malkin, Syed S. Islam, Karl F. Herold, Hugh C. Hemmings, Em B Sorum, Rie Nygaard, Paul M. Riegelhaupt

## Abstract

Establishment of electrical potentials across biological membranes is a universal feature of all cells. Tandem pore domain (K2P) potassium ion channels play pivotal roles in maintaining cellular membrane potentials, shaping physiological responses across a diverse range of cell types. With only a limited repertoire of high-aAinity and subtype-selective K2P modulators available for experimental or therapeutic use, we devised a strategy to genetically engineer K2P channels that are potently activated by rapamycin or non-immunomodulatory rapamycin analogs. Insertion of the FRB domain of mTOR into a short flexible cytoplasmic loop between the second and third transmembrane (TM) domains of the TREK1 K2P channel yielded fusion channels that are activated by nanomolar concentrations of rapamycin. Rapamycin-induced potentiation requires recruitment of an FKBP binding partner, from either the endogenous pool of FKBP within the cell or through fusion of FKBP to the C-terminus of TREK1. Formation of an FRB/rapamycin/FKBP ternary complex within the core of the TREK1 channel leads to an increase in TREK1 single-channel open probability and unitary current, mimicking positive modulatory eAects of conventional TREK1 activating cues. Cryo-EM structures demonstrate that rapamycin-induced ternary complex formation rigidifies the position of the FRB and stabilizes the TM2/TM3 loop in an active channel conformation. We demonstrate that FRB fusion can be employed to successfully activate several K2P channel isoforms, providing chemogenetically targetable tools for direct manipulation of cellular membrane potential.

## Introduction

Tandem pore domain (K2P) potassium channels play central roles in determining cellular resting membrane potentials, fine-tuning electrical excitability throughout the human nervous system. In excitable tissues, K2P channels contribute to the input resistance that determines electrical firing thresholds. Across non-excitable tissues, alterations in resting membrane potential influence functions such as smooth muscle contractility, endocrine hormone secretion, and rates of cell cycle progression, proliferation, and migration in healthy and cancerous cells^1^. Inheritable or acquired alterations in K2P channel activity leads to a spectrum of pathophysiological disease states^2,3^, highlighting the importance of this ion channel family in maintaining normal physiological homeostasis.

The various K2P subfamily members exhibit unique tissue distributions and distinctive sensitivities to physiological and pharmacological regulators^4^, but there are few high-affinity and subtype-selective small molecules that modulate individual K2P subfamily members. Certain K2P subtypes can be selectively inhibited by high-affinity small molecules or peptides^5–7^, but K2P channel activators have proven particularly difficult to develop. Despite multiple unbiased high-throughput screening efforts^8–12^, small-molecule activators have only micromolar affinity and are frequently promiscuous across K2P subtypes. Here, we describe the development of a set of genetically encodable engineered K2P channels that are potently activated by the macrolide small molecule rapamycin or non-immunomodulatory rapamycin analogs.

Rapamycin binds with high affinity to two discrete small protein domains: the peptidyl-prolyl-isomerase FK506 Binding Protein (FKBP) and the FKBP-rapamycin binding (FRB) domain of mTOR (mammalian Target of Rapamycin). While FKBP and FRB have no intrinsic affinity for each other, they form a tightly bound nanomolar-affinity ternary complex with rapamycin. This distinctive property has made FKBP/rapamycin/FRB a powerful protein engineering tool. Genetic fusion of FRB and FKBP to proteins of interest allows rapamycin-induced FKBP/FRB dimerization to effectively deliver proteins of interest to their site of action or to binding partners. Rapamycin-induced dimerization has found a wide range of applications, including control of gene expression^13^, modulation of membrane-associated lipid phosphatase^14^, kinase^15^, or GTPase activities^16,17^, and inducible control of transmembrane signal transduction^18^. While these examples utilize rapamycin as a vehicle to promote inducible protein targeting, rapamycin complex formation can also modulate intrinsic biophysical properties of the FRB and FKBP proteins. Degradation-prone mutants of FRB or FKBP are stabilized by rapamycin binding, such that fusion of the degradation-prone domains to proteins of interest allows for selective rapamycin-induced degradation or stabilization^19–21^. Rapamycin can also serve as an engineered enzymatic ligand, with fusion of FKBP to the catalytic domain of focal adhesion kinase (FAK) found to permit rapamycin to allosterically stimulate kinase activity^22^. Building on these approaches, we here show that the high-affinity interactions between rapamycin, FRB, and FKBP can be utilized to modulate K2P ion channel conformation and promote channel opening.

## Results

To engineer a rapamycin-responsive K2P channel, we introduced the FKBP-rapamycin-binding (FRB) domain of mTOR into the short intracellular loop that links the second (TM2) and third (TM3) transmembrane domains of the TREK1 channel (Figure 1a, b). The FRB N-terminus was fused to the TREK1 W199 residue at the end of TM2, while the C-terminus of FRB was joined to the TM3 Q203 residue, with deletion of three intervening residues from TREK1 (Figure 1a inset). Expression of this chimeric TREK1 M2/M3 FRB fusion construct (referred to as FRB0) in *Xenopus laevis* oocytes yielded a functional potassium channel that was indistinguishable from wild-type TREK1 (Figure 1 and Supplemental Figure 1). Insertion of FRB had no significant impact on TREK1 basal current density, gating by temperature, intracellular acidification, or membrane tension (Supplemental Figure 1a, b, g and Figure 4b, c). Sensitivities to the TREK1 activator BL1249 or inhibitor fluoxetine were similarly unperturbed by FRB insertion (Supplemental Figure 1c–f). As both fluoxetine and BL1249 bind to a fenestration site^23,24^ lined by the TM2 and TM3 helices that flank the FRB, we took the absence of changes in BL1249 or fluoxetine drug sensitivities as an indication that FRB insertion does not induce a significant local perturbation in TREK1 channel structure. However, unlike TREK1 WT channels, which are insensitive to the macrolide immunomodulatory drug rapamycin, the FRB0 channel was potently activated by rapamycin at mid-nanomolar concentrations (EC_50_ 55 nM, Figure 1c-e), exceeding the mid-micromolar potencies of BL1249^24–26^, ML335^27,28^, or any other conventional small molecule TREK1 activator^8,9,29–32^.

**Figure 1:**
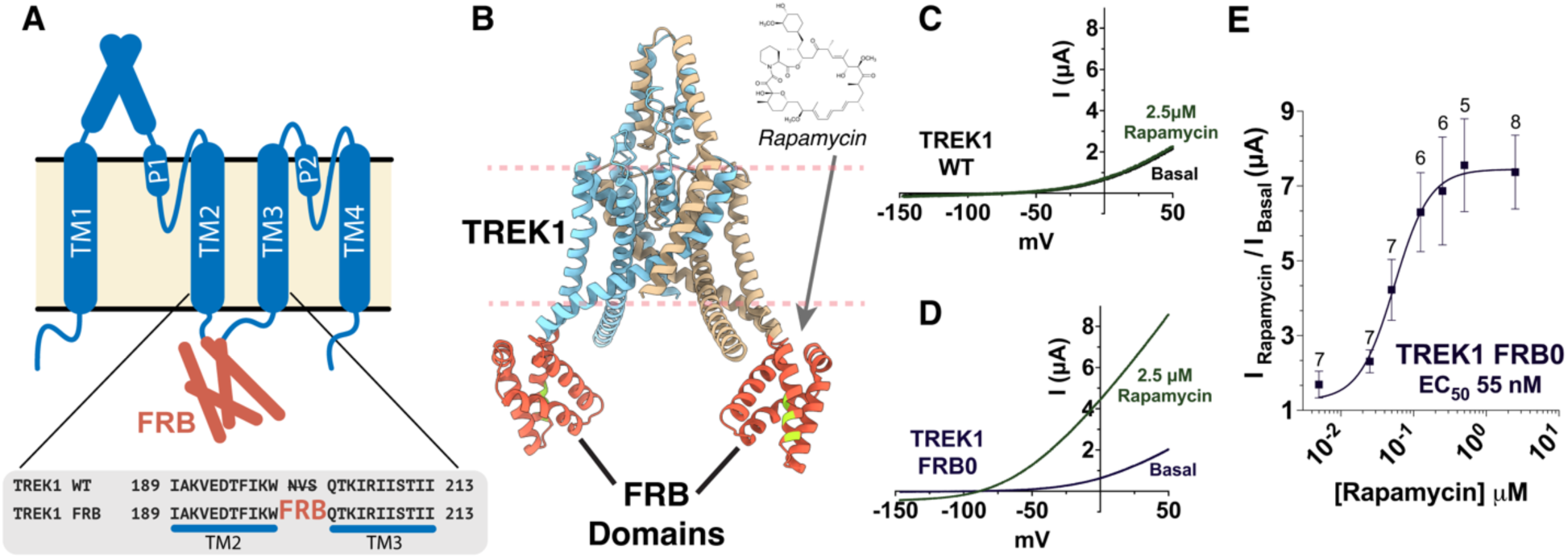
Designing a rapamycin responsive K2P channel. **A.** An illustration of the transmembrane topology of the TREK1/FRB fusion channel (FRB0), showing the position of the inserted FRB domain (orange) within the TREK1 TM2/TM3 loop. The protein sequence of the TREK1 distal TM2 and proximal TM3 domains is shown in the inset, with location of the inserted FRB protein sequence noted. **B.** An alphafold^75^ protein structure prediction for the FRB0 channel. The rapamycin binding interface within the FRB domain is shown in yellow. **C,D.** Representative TEVC recordings from *X. laevis* oocytes, showing TREK1 WT (C) or TREK1 FRB0 (D) currents elicited by a voltage ramp in the absence or presence of 2.5 µM rapamycin. **E.** Rapamycin dose response relationship for the FRB0 fusion channel. Number of replicates for each point is shown; error bars represent SEM.

### Activation by rapamycin requires recruitment of endogenous FKBP

The nanomolar potency of rapamycin for activation of the FRB0 channel was surprising, as a previous biophysical study of the interaction between rapamycin and the isolated FRB domain reported a much lower micromolar range K_d_^33^. Nanomolar affinity binding between rapamycin and FRB is only described when rapamycin is complexed with an FKBP protein. As we had not heterologously added FKBP to our experimental system, we explored whether endogenous *Xenopus laevis* oocyte FKBP^34^ contributes to FRB0 channel activation and the potent effect of rapamycin.

To test for involvement of endogenous FKBP, we utilized a Rapa*-3a analog of rapamycin^35^ that features a chemical modification designed to reduce the affinity of Rapa*-3a for FKBP (Figure 2a–c). Treatment of FRB0 with Rapa*-3a (Figure 2c, f) produced an equivalent degree of channel activation as observed with rapamycin, but at 8.7-fold lower potency (EC_50_ 480 nM). As the FRB binding interfaces of Rapa*-3a and rapamycin are identical (Figure 2b, c), the observed shift in Rapa*-3a potency suggests involvement of FKBP binding in the FRB0 channel response. To further demonstrate involvement of FKBP, we co-expressed FRB0 with soluble human FKBP12 bearing a compensatory V55G mutation (Figure 2a), designed to restore the binding affinity of Rapa*-3a for FKBP12^35^. This change in FKBP rescued the potency of Rapa*-3a to that of rapamycin (Supplemental Figure 2h).

**Figure 2:**
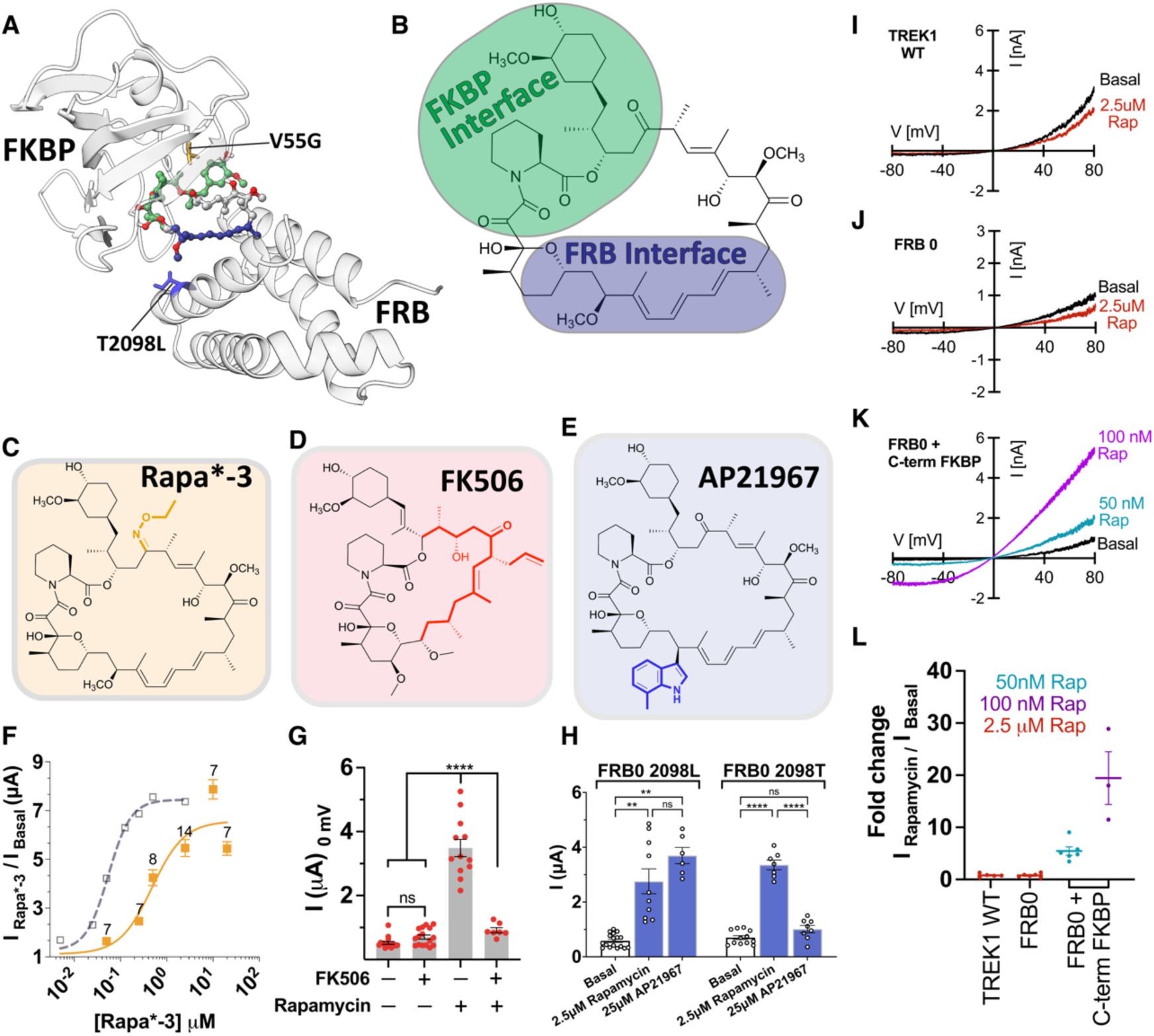
Rapamycin modulation requires FRB/FKBP/rapamycin ternary complex formation. **A.** A crystallographically derived model of the FRB/FKBP/rapamycin complex (PDB: 1NSG)^42^. The protein structures of FRB and FKBP are notated and shown in grey, while the rapamycin molecule is colored to illustrate the FKBP (orange) and FRB (purple) binding surfaces, to match **B.** a molecular representation of rapamycin **C, D, E.** Chemical structures of Rapa*-3, FK506, and AP21967, highlighting the substituted ethyloxime moiety present in Rapa*-3 (orange), the calcineurin binding interface of FK506 (red), or the substituted 7-methyl-indole moiety in AP21967 (blue). **F.** Dose response relationship of FRB0 channel activation by Rapa*-3, in the absence (solid orange squares) or presence (open orange triangles) of co-expressed FKBP V55G protein. Number of replicates for each point is shown; error bars represent SEM. Grey dashed curve represents dose response of the FRB0 channel for rapamycin. **G.** Current elicited from FRB0 channels in TEVC recordings from *X. laevis* oocytes, after treatment with FK506 or rapamycin **H.** Current elicited from FRB0 2098T or 2098L channels in TEVC recordings from *X. laevis* oocytes, after incubation with saturating doses of either rapamycin or AP21967 (AP21967 dose response curve shown in Supplemental Figure 2K). **I,J,K.** Inside-out patch clamp recordings from *X. laevis* oocyte membranes, for TREK1 WT (I), TREK1 FRB0 (J), and TREK1 FRB0 C-term FKBP (K) channels, prior to and following administration of rapamycin. **L.** Quantitation of rapamycin responses in inside-out patch clamp experiments.

We obtained complementary results utilizing FK506 (tacrolimus) as a competitive inhibitor for rapamycin binding to FKBP (Figure 2d, g). FK506 lacks the chemical interface to bind FRB and should therefore have no direct effect on FRB0-rapamycin binding. FK506 and rapamycin do, however, share identical FKBP-binding interfaces (Figure 2b, d). Treatment with FK506 alone had no effect on FRB0 channel currents (Supplementary Figure 2a), but pretreatment with FK506 and subsequent co-application of equimolar doses of FK506 and rapamycin blocked rapamycin-induced FRB0 activation (Figure 2g, Supplementary Figure 2b). This is consistent with FK506 competing with rapamycin for FKBP binding, preventing channel activation by blocking rapamycin’s ability to recruit FKBP to the channel. Binding of the rapamycin/FKBP complex to the TREK1 embedded FRB domain was also essential for FRB0 channel activation, as the rapamycin analog AP21967 bearing a modification at the FRB binding interface (Figure 2e) failed to activate FRB0 in the absence of a T2098L mutation designed to compensate for the AP21967 modification (Figure 2a, h). These results demonstrate that formation of an FRB-rapamycin-FKBP ternary complex within the TREK1 TM2/TM3 loop is required for rapamycin-induced activation of the FRB0 channel in oocytes.

Consistent with the role of endogenous *Xenopus laevis* FKBP protein in the FRB0 response, activation by rapamycin was lost in inside-out patch clamp experiments where the plasma membrane is isolated from the cytoplasmic milieu (Figure 2j). Fusion of human FKBP12 to the C-terminus of the FRB0 channel was sufficient to reconstitute FRB0 rapamycin responsiveness in inside-out patch clamp recordings (Figure 2k, l). Endogenous levels of FKBP in *Xenopus* oocytes appeared sufficient to fully activate the FRB0 channel in whole cell TEVC recordings, as rapamycin potency and fold activation were unaffected by fusion of human FKBP12 to the C-terminus of FRB0 (Supplementary Figure 4e). In mammalian cells, FKBP availability appeared to be more limiting. In whole-cell patch clamp recordings from HEK293T cells expressing FRB0, currents were clearly potentiated by rapamycin in the absence of exogenous FKBP, though enhancement of the rapamycin effect was observed after fusion of a C-terminal FKBP to the FRB0 channel (Supplemental Figure 3).

### Rapamycin binding promotes conventional TREK1 gating

To characterize the molecular basis for potentiation of FRB0 currents by rapamycin, we examined rapamycin’s influence on the single-channel properties of the fusion channel (Figure 3). In the absence of rapamycin, FRB0 channels exhibited a low open probability (P_o_ 0.002 ± 0.0005 SEM), with two kinetically distinct closed states featuring short (C_1_, 1.05 ms) or long (C_2_, 202 ms) dwell times (Figure 3a–c, upper panels). Application of rapamycin (Figure 3a–c, lower panels) eliminated the appearance of the long dwell time C_2_ closed state, promoting an overall increase in open channel probability (P_o_ 0.26 ± 0.07 SEM). Rapamycin also favored openings of a larger unitary current size (Figure 3e). Whereas most channel openings in the absence of rapamycin exhibited unitary currents of ∼1 pA, application of rapamycin resulted in the appearance of a ∼2.5 pA higher conductance state of the channel. The related mechanosensitive K2P channel TRAAK^36^ shows analogous patterns of single-channel behavior in response to activation by applied membrane tension^36^.

**Figure 3:**
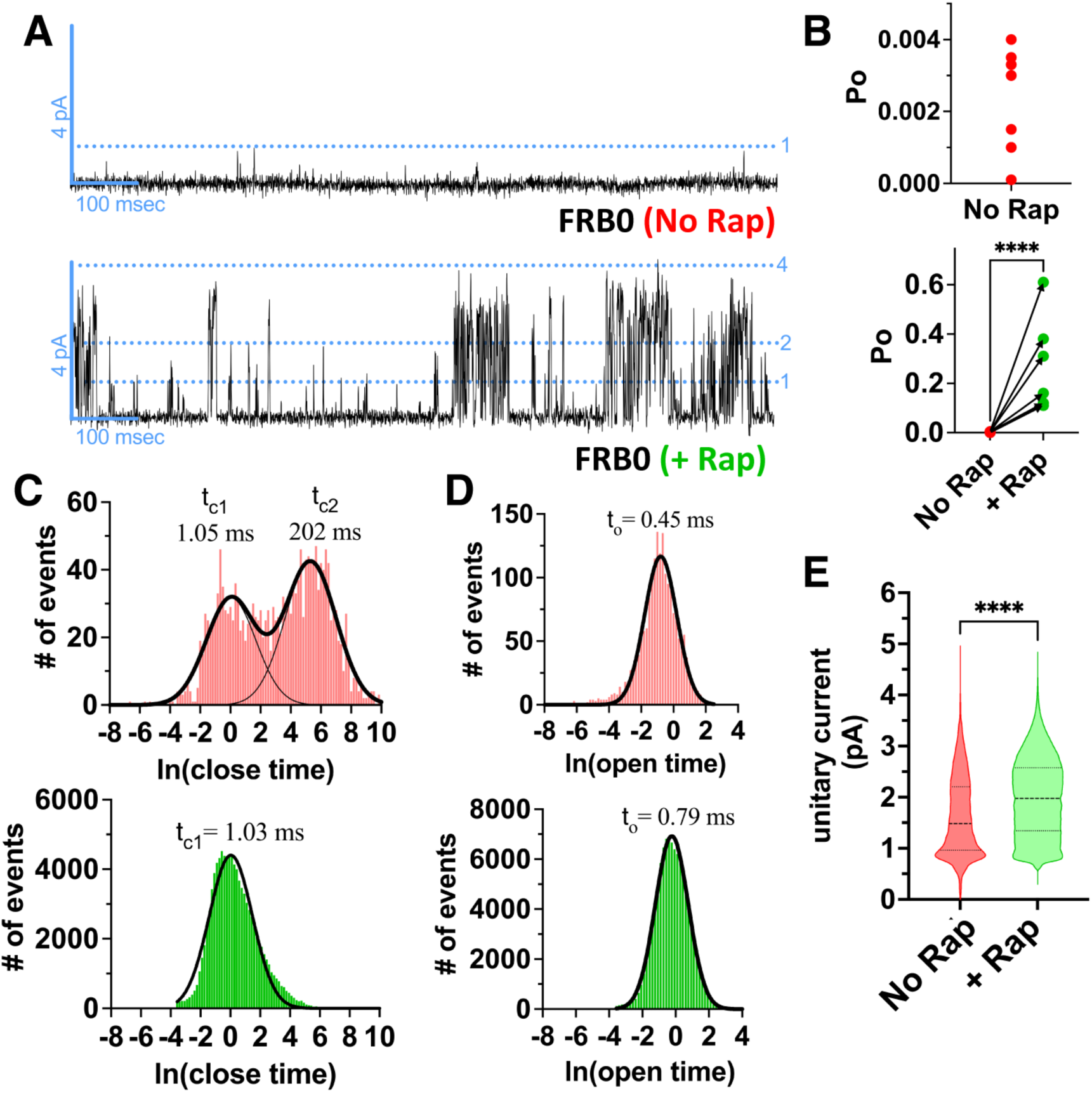
Effects of rapamycin on single-channel properties of the FRB0 channel. **A.** Representative single-channel recording from an inside-out patch of *Xenopus laevis* oocyte membrane expressing an FRB0 C-terminal FKBP channel, prior to (upper panel) and following application of 25 µM rapamycin (lower panel). **B.** Single channel open probability for 7 distinct recordings, prior to (upper panel, red) and following (lower panel, green) addition of rapamycin. Arrows demonstrate the degree of change in Po induced by rapamycin application. Statistical significance was determined by paired Student’s *t*-test, results indicated, ****p<0.0001. **C**, **D.** Distributions of (C) closed and (D) open dwell times, prior to (upper panels, red) and following application of 25 µM rapamycin (lower panel, green). **E.** Violin plots showing single channel unitary current from all channel openings, prior to (upper panels, red) and following application of 25 µM rapamycin (lower panel, green). Statistical significance was determined by Mann-Whitney test, results indicated, ****p<0.0001.

**Figure 4:**
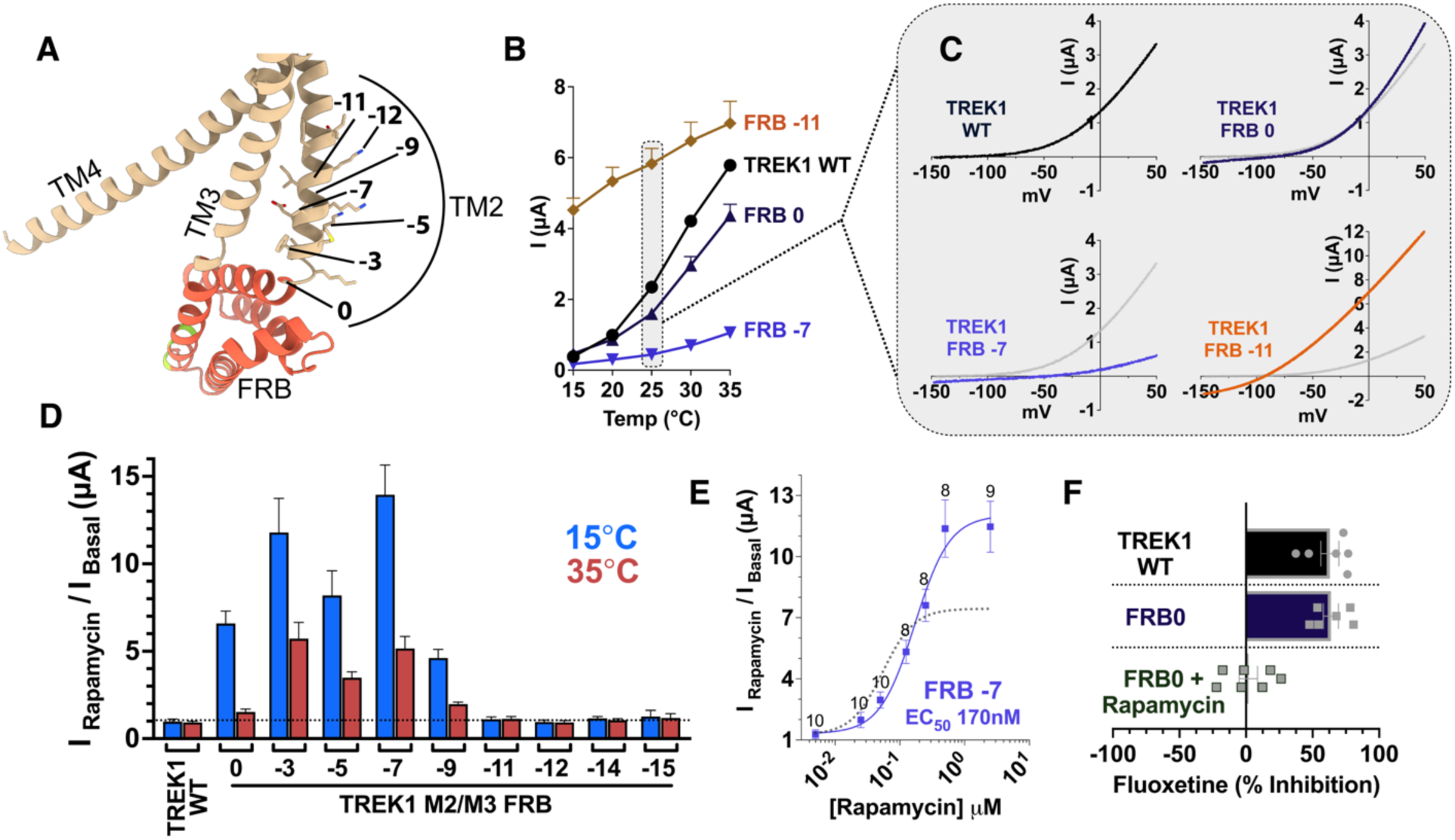
Altering the junction point between TREK1 and FRB modulates channel function. **A**. Zoomed in view of the FRB0 alphafold model, highlighting the insertion site for the FRB domain. Alternative TM2 junction sites for FRB insertion are noted. **B.** Temperature dependence of TREK1 WT or the indicated FRB fusion constructs, assayed by TEVC recordings of *Xenopus laevis* oocytes at temperatures from 15–35°C, in 5°C increments. **C**. Representative TEVC I/V curve traces assayed at 25°C. The TREK1 WT I/V curve (top left panel) is reproduced (grey) in all other panels for the purpose of comparison. **D.** Normalized fold change response to application of a saturating 2.5 µM dose of rapamycin in TREK1 WT or FRB fusion constructs, assayed in cold inactivated (15°C) or heat activated (35°C) conditions. **E.** Dose response relationship of FRB-7 channel activation by rapamycin. Number of replicates for each point is shown; error bars represent SEM. Grey dashed curve represents dose response of the FRB0 channel for rapamycin. **F.** Percent inhibition of channel currents by application of 200 µM Fluoxetine, in TREK1 WT, FRB0, or in FRB0 channels treated with 2.5 µM rapamycin.

Perturbations to residues within the distal TM2 region of the K2P architecture have been shown to potentiate mechanosensitive K2P channel activity^37^, and we hypothesized that binding of rapamycin/FKBP to the FRB0 channel might promote channel activation by inducing changes in the conformation or dynamics of the TM2/TM3 loop. To explore this possibility, we created multiple additional TREK1/FRB fusion constructs, altering the fusion point between the N-terminus of FRB and the TREK1 TM2. We progressively deleted residues from the TREK1 TM2 helix in sequentially numbered fusion constructs, such that an FRB-n construct corresponds to a TREK1 M2/M3 FRB fusion with n-number of residues before W199 deleted (Figure 4a).

All TREK1/FRB fusions from FRB0 to FRB-12 yielded functional channels, though the properties of these chimeric channels varied. The FRB-3 and FRB-7 constructs exhibited small basal currents that were minimally responsive to activation by heat, while the FRB-11 and FRB-12 channel constructs were highly active at room temperature and resistant to inhibition by cold (Figure 4b, c, and Supplemental Figure 5a, c). The FRB-5 and FRB-9 fusions exhibited an intermediate phenotype, with reduced temperature sensitivity due to both diminished inhibition by cold and limited activation by heat (Supplemental Figure 5b). Deletion of TM2 residues past K187 at the-13 position appeared to fully disrupt TREK1 channel integrity, as the FRB-14 and FRB-15 constructs exhibited no discernible TREK1 currents at any temperature (Supplemental Figure 5d, e).

For fusion constructs from FRB0 through FRB-9, application of rapamycin led to potentiation of TREK1 currents (Figure 4d). Rapamycin potentiation was most pronounced when basal currents were inactivated by cold (15°C), and the effects of heat activation and rapamycin potentiation were non-additive. This was most clearly exemplified in the FRB0 fusion construct that exhibits a temperature-gating profile similar to WT TREK1 (Figure 4b). Application of rapamycin to the FRB0 channel at 15°C produced a >7-fold increase in channel activity, whereas full activation of the FRB0 by heating to 35°C led to a complete loss of rapamycin potency (Figure 4d). Rapamycin was similarly ineffective in the proximal FRB fusion constructs FRB-11 and FRB-12, where the FRB insertion itself appeared to promote channel opening and mimic full heat activation (Figure 4d). Potentiation by rapamycin after heating to 35°C was only readily observed in the FRB-3 through FRB-9 constructs (Figure 4d), as the extent of TREK1 activation by heat was reduced in these fusion constructs and allowed for additional TREK1 activation upon rapamycin binding (Supplemental Figure 5b).

While alteration of the fusion point between TREK1 and FRB resulted in large changes in activation profiles for the resultant fusion channels, the affinity of rapamycin/FKBP for the FRB fragment appeared only minimally affected. In the FRB-7 channel that exhibited the most restricted temperature-activation profile, rapamycin potentiation exhibited an EC_50_ of 170 nM (Figure 4e). While this affinity is modestly less potent than that for the FRB0 channel (EC_50_ 55 nM; Figure 1e), the low basal activity level of the FRB-7 channel resulted in an enhanced 13-fold increase in current density after application of rapamycin (Figure 4e).

TREK1 channel activation by heat, membrane tension, and volatile anesthetics are all believed to involve translation of the TREK1 TM4 helix to an “up” facing position^36,38^, a conformational rearrangement that promotes an open pore and conductive selectivity filter. The observed increase in single-channel unitary current following rapamycin administration and the loss of rapamycin potentiation upon heat activation suggest that rapamycin binding induces gating movements that mimic those known to occur in response to conventional TREK1 gating cues. In support of this notion, we find that application of rapamycin to the FRB0 fusion construct eliminated TREK1 fluoxetine inhibition (Figure 4f, Supplemental Figure 1e, f), similar to the loss of fluoxetine sensitivity observed in heat-activated TREK channels^39^. Fluoxetine has been shown to bind at a fenestration formed when the TREK TM4 helix is positioned in the “down” conformation^23^ and becomes ineffective as an inhibitor when activating stimuli close this fenestration by shifting TM4 to the “up” conformation^39–41^. The loss of fluoxetine sensitivity after rapamycin treatment suggests that binding of the rapamycin/FKBP complex to the FRB fragment within the TM2/TM3 loop modulates the position of the conformationally flexible TREK1 TM4 helix, promoting the stimulus-activated TM4 “up” conformation to potentiate channel currents.

### Flexibility of FRB within the TM2/TM3 loop is restricted by rapamycin/FKBP binding

To explore the mechanism by which rapamycin-induced ternary complex formation leads to TREK1 channel opening, we determined cryo-EM structures of the TREK1 FRB0 fusion protein in the absence or presence of rapamycin and FKBP. FRB was sub-cloned into the M2/M3 loop of a biochemically tractable C-terminally truncated *Danio rerio* TREK1 construct (drTREK1 M2/M3 FRB0 C-term Δ321, FRB0_EM_). The FRB0_EM_ construct retained the rapamycin responsiveness of the full-length FRB0 channel (Figure 5c), and purification of the FRB0_EM_ construct was uncomplicated (Supplemental Figure 6a–c). Fluorescent size exclusion chromatography (FSEC) profiles of a mixture of purified FRB0_EM_ channel and YFP/FKBP fusion protein showed a rapamycin-induced peak shift consistent with complex formation (Figure 5a). By varying the rapamycin concentration, the ratio of FRB0_EM_-bound *versus* unbound YFP/FKBP protein indicated an affinity of association of 137 nM (Figure 5b), comparable to the 55 nM EC_50_ for FRB0 channel activation in electrophysiological recordings. These control experiments demonstrate that the FRB0_EM_ construct recapitulates all relevant rapamycin-induced behaviors of the full-length FRB0 channel.

**Figure 5:**
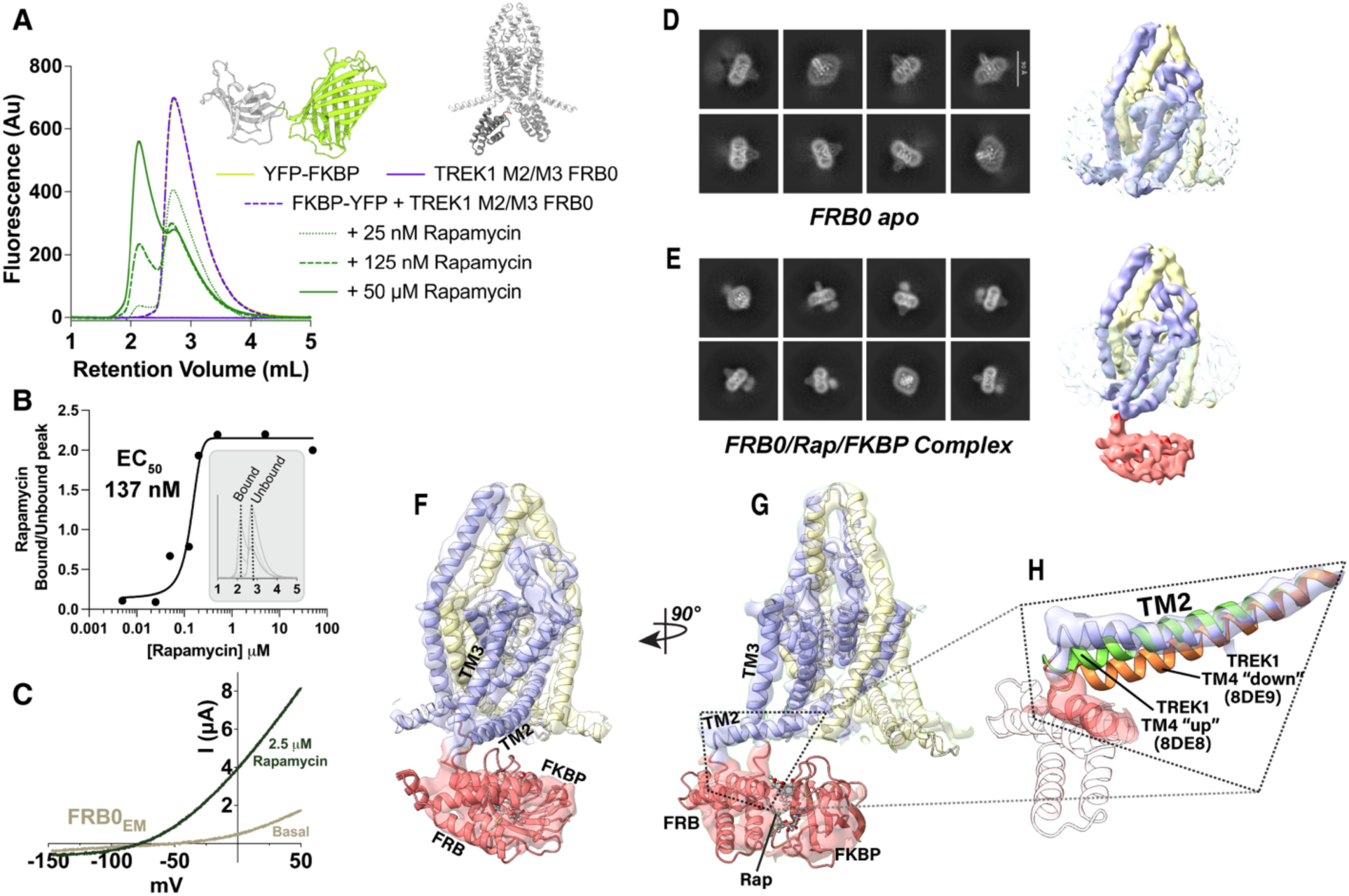
Cryo-EM structure of the FRB0 channel in complex with rapamycin and FKBP. **A**. Fluorescence size exclusion chromatography (FSEC) traces of purified YFP-FKBP (yellow), FRB0 channel (blue), or a 10:1 molar ratio mixture of YFP-FKBP to FRB0 in the absence (blue dashed) or presence (green) or varying concentrations of rapamycin. **B.** FSEC data from A, with the ratio of the fluorescence value for the bound versus unbound peaks (as shown in the grey inset) plotted on the Y-axis and rapamycin concentration plotted on the X-axis. **C**. Representative TEVC I/V curve traces assayed at 25°C, for the FRB0_EM_ construct, in the absence (tan) or presence of 2.5 µM rapamycin (brown). **D,E.** Representative 2D classes and a 3D reconstruction of the FRB0 apo sample (D) or the FRB0 channel in complex with rapamycin and FKBP (E). **F,G.** An overlay of the cryo-EM map density with a model of the FRB0 channel in complex with rapamycin and FKBP. **H.** Comparison of the position of the TM2 helix in the rapamycin and FKBP complexed FRB0 channel (blue) with the position of TM2 in previously solved structures of TM2 from a POPE bound and inactivated TREK1 TM4 “down” structure (pdbID 8DE9) or a POPA bound and activated TREK1 TM4 “up” structure (pdbID 8DE8).

We first determined the structure of the FRB0 channel in the absence of rapamycin or FKBP, obtaining a 4.33 Å intermediate-resolution map (Figure 5 and Supplementary Figure 7). In 2D classes and a 3D reconstruction (Figure 5d), the TREK1 transmembrane helices and cap domain were well resolved, but the FRB domains appeared flexible. The presence of the FRB was only apparent in 2D classes as a diffuse hazy density, and these flexible domains were absent in a 3D reconstruction of the channel (even at extended volume thresholds sufficient to visualize the surrounding detergent micelle, Figure 5d). Despite this flexibility, the influence of the inserted FRB domains was evident in the overall TREK1 conformation. Compared with a previously determined cryo-EM structure of WT TREK1^38^, FRB0_EM_ exhibits an upward movement of the proximal TM2 helix and loss of the structured density for the distal TM2/TM3 loop region that connects TREK1 to the FRB (Supplementary Figure 10a). Positioning of the TM2 helix is known to couple to TM4 movement in mechanosensitive K2Ps^37,38^, and in the FRB0_EM_ channel structure the TM4 gating helices are both in the active “up” conformation (Supplementary Figure 9b), differing from the apo WT TREK1 structure^38^ where the two TM4s are positioned asymmetrically in a mixed “up/down” conformation. These structural differences suggest a conformational path by which rapamycin-induced movement of TM2 and TM4 might ultimately activate TREK1, though we note that in the absence of rapamycin the functional properties of the TREK1 FRB0 channel are indistinguishable from TREK1 WT (Supplementary Figure 1 and Figure 4b). We hypothesize that the unrestricted mobility of the un-complexed FRB domains within the TM2/TM3 loop may preclude any functional impact of FRB insertion prior to the addition of rapamycin.

Vitrification of the FRB0_EM_ protein in rapamycin alone resulted in 2D classes with similarly dynamic FRB domains (Supplementary Figure 6d), but ternary complex formation with rapamycin and FKBP rigidified the positioning of the FRB within the TM2/TM3 loop. In a 4.2 Å reconstruction of the FRB0 channel complexed to rapamycin and FKBP (Figure 5e), the static position of the ternary complex was evident in both 2D classes and a 3D reconstruction. We docked a previously determined crystallographic structure of the FRB/rapamycin/FKBP ternary complex^42^ into the 3D reconstruction and could unambiguously fit the FRB helices into the experimentally determined density (Figure 5f). Surprisingly, despite a sufficient supply of rapamycin and FKBP for maximal occupancy of the FRBs, we only ever observed a single-bound FKBP. Modeling of two FRB/FKBP complexes at the cytoplasmic face of the channel (Supplementary Figure 6e) results in a steric clash between the bound FKBPs that may prevent complex formation on both sides of the TREK1 channel.

Preliminary AlphaFold modeling of the TREK1/FRB fusion (Figure 1b and Supplementary Figure 9a) suggested that the FRB domains would extend downward toward the cytoplasm, but in our cryo-EM structure, the FKBP-complexed FRB domain folds directly below the TM2 helix (Supplementary Figure 9a). Well-resolved density for the full span of the TREK1 TM2 helix as it connects to the FRB domain (Figure 5h) demonstrates that the membrane-proximal positioning of FRB promotes an upward translation of the TM2 helix, the same conformational rearrangement we observed in the un-complexed FRB0_EM_ structure (Supplementary Figure 10a, b). Further processing of a subset of the particles from the FRB/rapamycin/FKBP dataset where the density for the FRB/FKBP complex was absent (Supplementary Figure 8c), produced two additional higher-resolution reconstructions of TREK1 in TM4 “up” (3.64 Å) or asymmetric TM4 “up/down” (3.74 Å) conformations. We observe identical upward repositioning of the distal TM2 helices in these higher resolution reconstructions (Supplementary Figure 10c, d), indicating that upward translation of TM2 is a direct consequence of FRB fusion and not dependent on either TM4 position or rigid complexation with FKBP. This FRB-induced TM2 movement is an exaggerated version of the TM2 positioning found in the stimulus-activated TM4 “up” conformation of WT TREK1 channels^38^ (Figure 5h), though our results suggest that structural coupling between TM2 and TM4 may be altered by FRB insertion. We find that the effect of FRB insertion on TM2 position is present even in a subunit where TM4 is in the “down” position (Supplemental Figure 10d) and in the structure of the FRB0_EM_ channel with a rigidified FRB/FKBP complex (Figure 5d), the TM4 helix in the FRB fused subunit exhibits a mixture of TM4 “up” and “down” states (Supplementary Figure 9c). This ambiguous positioning of TM4 may reflect our inability to further sort the small number of particles in the reconstruction of the rigidified FRB/FKBP complex without losing resolution for the rigidified FRB/rapamycin/FKBP complex. Nonetheless, the sum of our structural results and functional data demonstrating that rapamycin-induced activation promotes channel properties consistent with the TM4 “up” conformation support a model in which rapamycin-induced rigidification of FRB position within the TM2/TM3 loop stabilizes the induced movement of TM2 to bias the channel toward a high activity TM4 “up” state.

### FRB fusion in other K2P channel family members

To test the generalizability of the FRB/rapamycin-based approach in other K2P family members, we inserted the FRB domain at equivalent sites within the TM2/TM3 loops of the closely related mechanosensitive TREK2 and TRAAK K2P channels (Figure 6a, b). In both cases, insertion of FRB within the TM2/TM3 loop yielded rapamycin-sensitive channels. Unlike the FRB0 channel that phenocopies WT TREK1 in the absence of rapamycin, both the TRAAK/FRB and TREK2/FRB fusion channels exhibited much smaller basal currents than those observed in the unmodified WT channels, and both showed a restricted responsiveness to activation by heat (Figure 6d, e). Despite this low basal activity level, both TRAAK/FRB and TREK2/FRB could be activated by application of rapamycin (Figure 6d,e), with the TRAAK/FRB channel exhibiting a rapamycin EC_50_ for activation of 38 nM (Supplementary Figure 9f), equivalent to that obtained for TREK1/FRB (Figure 1e).

**Figure 6:**
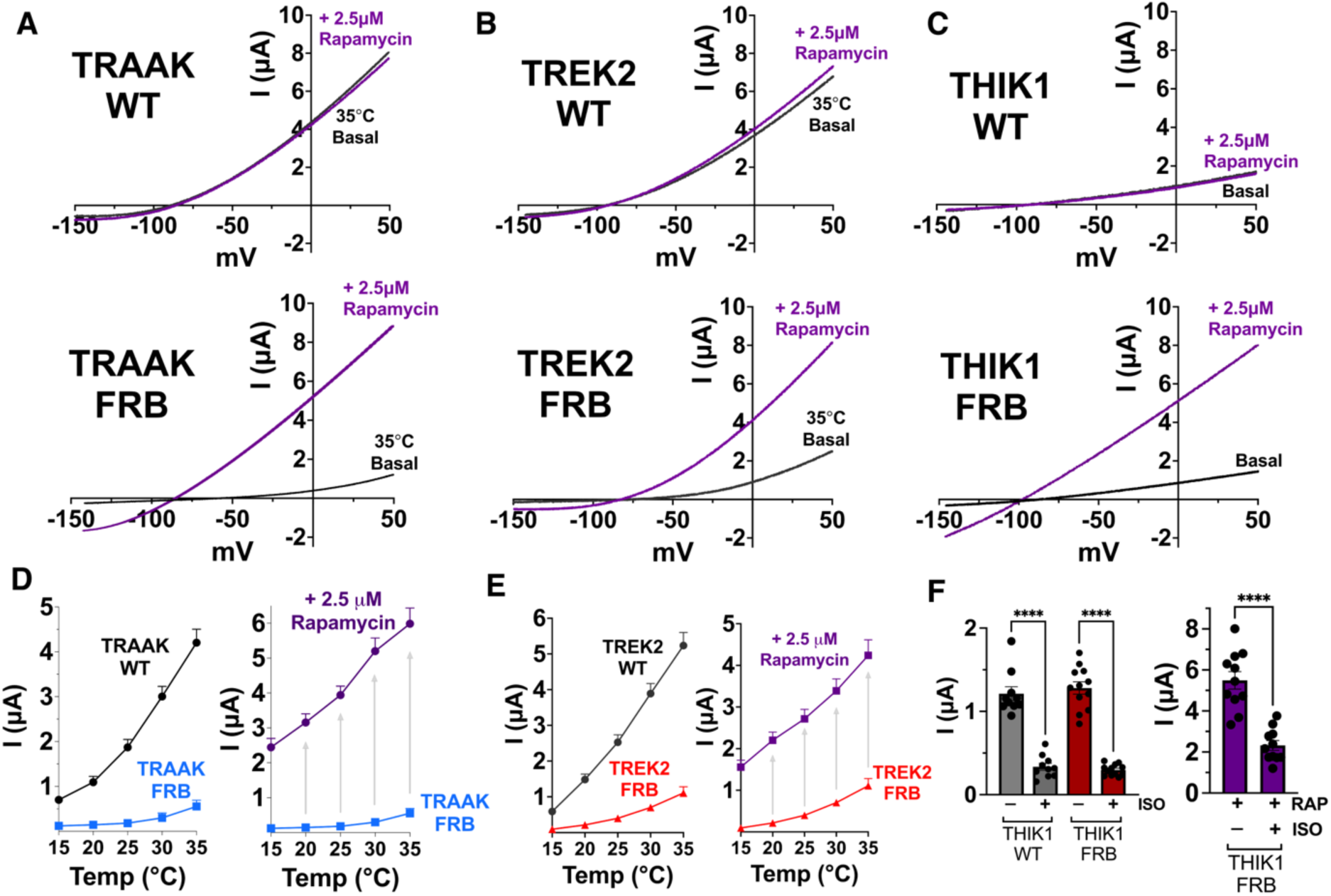
FRB fusion is effective in multiple alternative K2P channels. A,B,C. Representative TEVC I/V curve traces in the absence and presence of 2.5 µM rapamycin, from wildtype full-length human K2P isoforms (upper panels) or from the TM2/TM3 FRB fused chimeras (lower panels), for (A) TRAAK, (B) TREK2, or (C) THIK1. **D,E.** Temperature dependence of WT versus FRB chimeric (D) TRAAK or (E) TREK2 channels, assayed by TEVC recordings of *Xenopus laevis* oocytes at temperatures from 15–35°C, in 5°C increments (left panels). Current response to addition of 2.5 µM rapamycin at all temperatures is shown in right panels. **F.** Inhibition of THIK1 WT or THIK1 TM2/TM3 FRB chimeric channel currents after administration of 2 mM isoflurane (ISO) and **G.** isoflurane inhibition of the THIK1 TM2/TM3 FRB chimeric channel after administration of 2.5 µM rapamycin.

We then tested the generalizability of the FRB fusion approach in a K2P outside the mechanosensitive subfamily using THIK1, a K2P for which no small-molecule activators are available. THIK1 features a structured domain within the TM2/TM3 loop that plays an important role in channel gating^43^, and we fused the FRB domain between the end of the THIK1 TM2 and the beginning of this structured section of the TM2/TM3 loop. Insertion of the FRB at this site had no discernible impact on basal current size or on the inhibitory effect of the volatile anesthetic isoflurane (Figure 6f). The THIK1 binding site for isoflurane is positioned near the TM2/TM3 loop^43^, so the retained sensitivity to isoflurane in the THIK1/FRB fusion channel indicates that FRB insertion does not significantly perturb THIK1 channel structure. Like the mechanosensitive K2P/FRB fusion channels, the THIK1/FRB channel was potently activated by rapamycin (Figure 6c). Isoflurane inhibition of THIK1/FRB was diminished by rapamycin, similar to the effect of gain-of-function mutations within the anesthetic binding site near the TM2/TM3 loop region of THIK1^43^.

## Discussion

Studies of K2P channel gating frequently focus on the potassium selectivity filter^44–47^ or conformationally flexible pore-lining TM4 helix^37,38,48^, regions of the K2P architecture that mirror functionally important gating domains relevant to all potassium channels^49,50^. Considerably less attention has been paid to the TM2/TM3 loop region, a K2P-specific motif formed by the unique transmembrane topology of the tandem pore domain subfamily. Several lines of evidence highlight functionally important roles played by the TM2/TM3 loop region, including the presence of a modulatory volatile anesthetic binding site within this region^43,51–53^, gain-of-function mutations identified within the TM2 and TM3 helices of several K2Ps^37,43,54^, and structural evidence of coupling between the conformational movements of the TM2/TM3 loop and the functionally critical TM4 helix^37,38^. Our efforts to embed an FRB domain within the TREK1 TM2/TM3 loop were inspired by the presence of a channel modulatory calcineurin-binding domain within an extended TM2/TM3 loop in the TRESK K2P channel^55,56^. We show that rapamycin activation of our chimeric TREK1/FRB channel requires recruitment of FKBP, stabilizing the position of the flexibly embedded FRB domain and anchoring the TM2/TM3 loop in a position that favors TREK1 activation. A similar rapamycin-induced-rigidification mechanism is proposed to underlie rapamycin enhancement of FKBP-fused focal adhesion kinase activity^22^, such that our results raise the possibility for FRB (or FKBP) fusion to be a broadly generalizable means of engineering inducible control over protein conformation and function. Consistent with this notion, we show that multiple alternative K2P subfamily members can be fused to FRB and gated by rapamycin. We have not yet tested whether FRB insertion is effective across all K2P channel subfamilies, though recently published cryo-EM structures of channels from the TASK^57,58^, TALK^59^, and TWIK^7,60^ subfamilies should aid in the design of FRB insertions for these K2Ps.

Chemical genetic control of K2P channel activity provides a valuable platform for the study of K2P physiology *in-vivo*. The small size of the FRB (279 base pairs) and FKBP (322 base pairs) domains makes them readily amenable to CRISPR-targeted homologous directed repair (HDR) based knock-in approaches^61,62^, allowing for targeted insertion of these domains into K2P loci of immortalized cell lines or experimental animal genomes. As both the TREK1 FRB0 and THIK1 FRB channels exhibit normal channel gating and pharmacological properties in the absence of rapamycin (Figures 3, 6, and Supplemental Figure 1), insertion of FRB at these K2P loci should be well tolerated and functionally silent. These engineered FRB-fused TREK1 or THIK1 channels can then be activated by rapamycin or non-immunomodulatory rapamycin analogs within their native physiologic environment, avoiding confounding effects from heterologous channel over-expression or use of poorly selective small-molecule K2P activators. Similar functionally silent FRB insertion in the TREK2 or TRAAK channel will require additional design effort, as our initial FRB fusions in these backgrounds produce channels that exhibit diminished basal activity and restricted activation by heating (Figure 6d, e). As slight variations in the FRB fusion point in TREK1 produce variants that exhibit similarly low levels of basal activity and restricted heat sensitivity (Figure 4b, FRB-7), we anticipate that small alterations in the FRB fusion point in the TREK2 and TRAAK backgrounds are likely to provide functionally silent variants for these channels.

K2Ps modulate resting membrane potential and shape cellular electrical excitability, providing an opportunity for our rapamycin-responsive channels to also be deployed as chemogenetic tools to inducibly hyperpolarize cellular resting membrane potential and silence electrical activity. For this type of application, the K2P/FRB chimeras that exhibit very low intrinsic basal activity (including TREK1 FRB-7, TREK2 FRB, and TRAAK FRB) are ideal, as virus-mediated transgenic delivery of these low-activity K2P/FRB fusions should have minimal impact on cellular physiology prior to stimulation with rapamycin. Tissue or cell-type-restricted viral targeting of the K2P/FRB fusions should allow for systemic delivery of rapamycin or non-immunomodulatory rapamycin analogs to act on a narrow population of targeted cells. Additionally, the availability of rapamycin analogs that require photo-uncaging to permit FKBP binding^63^ may broaden the experimental utility of this approach. Photo-caged rapalogs can enable precise spatially targetable activation of K2Ps and may significantly improve the kinetics of rapamycin-based K2P activation, as the unmodified FRB binding interface in the photocaged derivatives should still complex with chimeric K2P/FRB channels even prior to light exposure. With the orders of magnitude larger single channel conductance of K2P channels relative to traditional opsin based optogenetic tools (TREK1 single channel conductance of ∼80 pS^64^ versus Channelrhodopsin-2 single channel conductance of ∼40 fS^65^), the non-inactivating nature of K2P channel currents, and the need for light stimulus only to produce uncaging, photo-caged rapamycin analogs may enable K2P/FRB chimeras to perform as versatile optogenetic tools to silence electrically active cells. CATKLAMP, an orthogonal bioengineering approach that uses derivatives of small-molecule K2P channel activators as chemogenetic tools to modulate mechanosensitive K2P activity, has already been shown to effectively silence the activity of hippocampal neurons^28^.

Several additional factors make rapamycin an ideal engineered K2P channel ligand. It is an FDA-approved, inexpensive, and commercially available drug that binds with high affinity and exhibits broad tissue distribution, including penetrance into the central and peripheral nervous system. Rapamycin’s one clear disadvantage as a protein engineering tool is its endogenous immunomodulatory properties. These arise from recruitment of FKBP to the FRB domain within the mammalian Target of Rapamycin (mTOR), with resultant inhibition of mTOR’s serine/threonine kinase activity leading to changes in cellular growth, metabolism, protein synthesis, and turnover^66^. Several groups have developed rapamycin analogs (rapalogs) that minimize or eliminate mTOR modulation but maintain the high-affinity interactions between rapalog and the FKBP and FRB domains^67^. We show that AP21967 and Rapa*3a, two such rapalog compounds, can be deployed as effective modulators of the TREK1/FRB fusion channel (Figure 2). For the case of AP21967, avoiding mTOR inhibition does come at the cost of lower affinity (Supplementary Figure 2k, 8.3 µM for AP21967 versus 55 nM for rapamycin) and slower kinetics, presumably due to reduced membrane permeability (Supplementary Figure 2c, half maximal effect 3 min for Rapamycin versus 18 min for AP21967). We note that the relatively lower potency of AP21967 still matches the highest-affinity TREK1-activating compounds presently available and retains the advantage of being selective for only genetically modified TREK1 constructs bearing the FRB T2098L domain. The Rapa*3a compound retains both the high affinity and rapid kinetics of rapamycin when FRB0 is expressed with FKBP V55G protein, but Rapa*3a was specifically designed to permit tissue-restricted mTOR activation at sites where FKBP V55G is expressed^35^. We show that Rapa*3a remains an effective TREK1 activator when FKBP V55G is fused to the C-terminus of the FRB0 channel (Supplementary Figure 2h), and tethering of the mutant FKBP to the membrane embedded channel should heavily favor association of Rapa*3a with the K2P-linked FKBP and FRB domains to minimize off-target interaction with mTOR.

The ternary complex formed by FRB, FKBP, and rapamycin is known to dissociate at an extremely slow rate^68^, such that rapamycin acts essentially as an irreversible activator of the K2P/FRB channels. While this property may be desirable for certain experimental paradigms, a more modular and graded switch to control K2P channel activity could provide additional versatility to the engineered design. Our results across several K2Ps show that the TM2/TM3 loop is permissive to insertion of exogenous domains, and we envision future creation of additional novel K2P tools with conformationally switchable protein domains inserted into the TM2/TM3 loop, creating tunable K2P rheostats for graded control of membrane potential.

## Methods

### Cloning and reagents

For insertion of FRB into K2P channel sequences, we used sequence and ligation-independent cloning (SLIC), with Phusion DNA polymerase (New England Biolabs) used for PCR reactions and T4 DNA polymerase treatment (in the absence of nucleotides) used to generate sticky ends. T4 treatment was terminated by addition of 1 mM dCTP (New England Biolabs). Insert and vector DNA were combined in T4 ligase buffer (New England Biolabs) and transformed into DH5α competent cells. For the initial TREK1/FRB fusion (FRB0), we inserted the FRB region of human mTOR (from residues 2021 to 2118, UniProt P42345) bearing the T2098L mutation into the TM2/TM3 loop of the *Mus musculus* TREK1 gene (mTREK1, K2P2.1, UniProt P97438), between residues W199 and Q203, with deletion of three intervening residues. The FRB-3 through FRB-15 variants of this original construct were subsequently created using QuikChange mutagenesis. SLIC cloning was used to add human FKBP12 (UniProt Q3V793) to the C-terminus of the FRB0 construct. For TREK2/FRB fusion, FRB was inserted into the *Homo sapiens* TREK2 gene (hTREK2, K2P10.1, UniProt P57789), between residues K229 and Q233, with deletion of the three intervening residues. For TRAAK/FRB fusion, FRB was inserted into the *Homo sapiens* TRAAK gene (hTRAAK, K2P4.1, UniProt Q9NYG8), between residues W186 and P189, with deletion of the two intervening residues. For THIK1/FRB fusion, FRB was inserted into the *Homo sapiens* THIK1 gene (THIK1, K2P13.1, UniProt Q9HB14), between residues G169 and W190, with deletion of twenty intervening residues.

Rapamycin was obtained from Santa Cruz Biotechnology (Dallas, TX). Rapa*3 (a mixture of the E and Z isomers), FK506, fluoxetine, and BL1249 were all obtained from Sigma-Aldrich. If not specified, all other chemical reagents were purchased from Sigma-Aldrich.

### Macroscopic electrophysiological recordings in Xenopus laevis oocytes

For electrophysiological studies in *Xenopus laevis* oocytes, wild-type full-length K2P genes or K2P/FRB chimeric fusion constructs were cloned into a pGEMHE plasmid (for TREK1) of a pfAW plasmid (for TREK2, TRAAK, and THIK1). K2P sequences were inserted downstream of the T7 RNA polymerase promoter region to facilitate in-vitro production of 5’ capped RNA (cRNA). The plasmids were linearized with FastDigest AflII (for pGEMHE) or MluI (for pfAW) restriction enzymes (ThermoScientific), purified using a QIAquick Nucleotide Removal Kit (Qiagen), and cRNA was produced using an mMessage mMachine Kit (T7 promoter, Ambion, Life Technologies). The products of the cRNA reaction were treated with DNAse, purified using an RNeasy RNA cleanup kit (Qiagen), and stored at −80°C prior to injection into oocytes. For all recordings, cRNA was injected into defolliculated stage IV-V Xenopus *laevis* oocytes (Xenoocyte, Dexter, MI), and the injected cells were stored in 1x Leibovitz’s L15 media (supplemented with 5 mM HEPES and 2.5% penicillin/streptomycin) at 16°C prior to recording.

For two-electrode voltage clamp (TEVC) studies, oocytes were injected with 2.5 ng of cRNA, and recordings were performed 24 – 36 hrs. after microinjection. Borosilicate microelectrodes were pulled to a resistance of 0.2 – 2.0 MΩ and backfilled with 3 M KCl solution. Oocytes were perfused with ND96 solution (96 mM NaCl, 2 mM KCl, 1.8 mM CaCl_2_, 2.0 mM MgCl2, 5 mM HEPES; pH 7.4) at a rate of 2.5 mL/min. Currents were acquired using an OC-725C Oocyte Clamp amplifier (Warner Instruments), controlled by pClamp10 software (Molecular Devices). A Digidata 1332A digitizer (MDS Analytical Technologies) was used to digitize the signal at 2 kHz. Currents were evoked from a −80 mV holding potential by a 500 ms ramp from −150 mV to +50 mV, with a 10 s wait time between each subsequent sweep. The temperature of the perfused solution was monitored using a CL-100-controlled thermistor placed in the bath solution immediately downstream of the oocyte and controlled by a SC-20 in-line heater/cooler combined with an LCS-1 liquid cooling system operated by a CL-100 bipolar temperature controller (Warner Instruments). For temperature experiments, the perfusate was warmed from 15°C to 35°C in 5°C increments, with recordings performed once temperature readings stabilized at the desired values. If not stated otherwise, recordings were carried out at room temperature (21°C). All TEVC data presented in the manuscript were obtained from at least two independent batches of ovaries, with a minimum of n = 5 experiments per condition.

For isoflurane experiments, a 99% isoflurane solution (Baxter Healthcare Corporation) was added to 100 mL of ND96 until a clear phase separation was achieved. The mixture was sealed tightly in a glass bottle and stirred vigorously overnight prior to usage. The maximal solubility of isoflurane in salt solution at room temperature has been empirically determined to be 15 mM^69^, and 2 mM isoflurane working concentrations were achieved by appropriate dilution of the aqueous phase of the saturated solution into ND96, using sealed syringes to prevent environmental loss of anesthetic.

Initial attempts to perform rapamycin treatment directly on our two-electrode voltage-clamp electrophysiology rig proved unreliable, as rapamycin appeared to bind tightly to both the perfusion tubing and the recording chamber on our rig and could not easily be removed between experimental trials. We found that rapamycin pre-treatment of oocytes in 6-well untreated culture dishes prior to electrophysiological studies was a far more reproducible approach. Immediately prior to recordings, injected oocytes were incubated at room temperature in drug-supplemented ND96 solution for 10 minutes (Rapamycin), 20 minutes (Rapa*), or 40 minutes (AP21967), with these individualized incubation times set by the plateau in peak drug effect, as determined in time course assays (Supplemental Figure 2c). Oocytes were washed 5X in an excess of ND96 buffer before application to the electrophysiology rig, to prevent carryover of rapamycin to the recording chamber.

For inside-out patch clamp studies, oocytes were injected with 1.25 – 2.5 ng of cRNA, and recordings were performed 48 – 72 h after microinjection. The bath electrode was installed via a KCL-agar salt bridge, and the reference electrode was pulled from thick-walled, fire-polished borosilicate glass pipettes (Sutter Instrument Co., #B200-116-10) using a Sutter Instrument Co. pipette puller Model P-97. Pipettes were polished using a microforge (MF2, Narishige Scientific Instrument, Japan) to achieve resistances ranging from 0.4 – 0.8 MΩ. Currents were measured using an EPC10 USB amplifier (HEKA electronics, DE), with a sampling rate of 10 kHz and data filtering at 2.9 kHz. Recordings were performed using either a ramp protocol (−80 to +80 mV, 1 s duration, 5 s pulse intervals) or a rectangle step protocol from −100 to +100 mV (50 ms duration) in 20 mV increments. Recordings were carried out in symmetrical potassium concentrations (120 mM), with the patch pipette filled with 120 mM KCl, 10 mM HEPES, and 3.6 mM CaCl_2_, adjusted to pH 7.4 with KOH/HCl and the bath solution containing 120 mM KCl, 10 mM HEPES, 2 mM EGTA, and 1 mM pyrophosphate adjusted to pH 7.4 (for measurement of basal current) or pH 5.5 (to measure channel activation). Pressure pulses were generated using an HSPC-2-SB pressure clamp (ALA scientific) installed to the side-port of the patch pipette. For rapamycin treatment, we installed a custom 3D-printed bath that was easily removable for cleaning and allowed for direct application of rapamycin to oocyte patches. Rapamycin-containing solution was manually added to the bath to achieve the desired concentration of drug.

### Single-channel recordings in Xenopus laevis oocytes

For single-channel recordings, currents were recorded at 25°C from inside-out patches excised from oocytes 1 days after injection with 0.25 ng of mTREK1 M2/M3 FRB C-term FKBP cRNA. Pipette solution contained (in mM) 15 KCl, 135 NaCl, 10 HEPES, pH = 7.4 with NaOH. The solution in the bath contained 150 KCl, 10 HEPES, 1 EGTA, pH = 7.1 with KOH. All single-channel records were made at a holding potential of 0 mV and are presented in physiological convention. To maintain the integrity of brief TREK1 dwell times, records were not further filtered. Baseline correction was performed manually based on the observable closed state for each recording; slight baseline drifts after solution exchange in our experiments were corrected. Portions of records from prior to application of rapamycin were analyzed separately from records after rapamycin application.

For dwell time analysis, single-channel open-close transitions were idealized by half-amplitude threshold crossing and dwell time event lists were generated for each baseline-corrected record. Dwell time histograms were generated from the dwell time event lists using custom-built software^70^; the histogram fits were generated in GraphPad Prism (v10.6.1) using single and double Gaussian fits. For unitary current analysis, event lists were first generated in Clampfit 10.7, applying a half-amplitude detection cutoff. Currents were filtered to 10 kHz and were analyzed with a custom script in Python 3.7 using the pyABF module^71^. Each point in the filtered record was annotated as closed or open using the events list. Violin plots were generated in GraphPad Prism (v10.6.1) from all individual unitary current measurements entered as raw values in column format. The violin shape represents the kernel density estimate of the full distribution of unitary current amplitudes. The mean was overlaid with the standard deviation (SD). The minimum and maximum unitary current values are indicated by the full vertical extent of the distribution.

### Electrophysiological recordings in HEK293 mammalian cells

HEK293T cells (CRL-3216 from ATCC, Manassas, VA) were maintained in high-glucose Dulbecco’s modified Eagle medium supplemented with 10% (vol/vol) fetal bovine serum (FBS), 2 mM GlutaMAX (Life Sciences, Waltham, MA) and 1% penicillin-streptomycin (Life Sciences). Passage numbers between 4 and 30 were used. Cells were seeded into 35-mm dishes and transfected with the respective TREK1 cDNA after 1–3 days using Lipofectamine 2000 (Life Sciences) according to the manufacturer’s protocol. One day after transfection, cells were released with TrypsinLE (Life Sciences) and replated at lower density onto 12-mm round #1.5 glass coverslips (Warner Instruments), a minimum of 3 h before recording isolated adherent cells with eGFP fluorescence.

Pipettes were pulled from standard borosilicate glass (1.5 mm OD/0.86 mm ID; Sutter Instrument) to a resistance of 1.5–2.5 MΩ using a P-1000 puller (Sutter Instrument). Whole-cell voltage-clamp electrophysiology was performed using an AxoPatch 200B amplifier (Molecular Devices) connected to a DigiData 1550B analogue-to-digital converter (Molecular Devices). Signals were sampled at 20 kHz and filtered at 5 kHz, respectively. Series resistance was corrected 85–90%.

Capacitive current transients were cancelled by the internal amplifier circuitry, and leak currents were subtracted using a standard P/4 protocol applied after the desired stimulus. Transfected cells were continuously perfused with extracellular solution at room temperature (24°C) containing (mM) 150 NaCl, 10 HEPES, 1 CaCl_2,_ 5 KCl, and 3 MgCl_2_, adjusted to pH 7.4 with NaOH. Pipette solutions contained (mM) 155 KCl, 3 MgCl_2_, 10 HEPES, 5 ethylene glycol tetraacetic acid (EGTA), adjusted to pH 7.20 with KOH. The osmolality of all solutions was balanced to 300±3 mOsm/kg H_2_O with sucrose.

Rapamycin (Santa Cruz) stock solutions were diluted to the desired working concentrations. Control solutions contained the same concentration of solvent (EtOH) as rapamycin solutions and did not exceed 0.048%, avoiding any bilayer-modifying effects. Solutions were delivered using a pressurized perfusion system (ALA Scientific Instruments) with Teflon tubing via a 200-μm diameter, low-dead-space manifold tip positioned ∼200–300 μm from the recorded cell. After control recordings, rapamycin was perfused before subsequent recordings and continuously thereafter. The perfusion chamber was rinsed with 80% isopropanol followed by distilled water in between experiments to remove any traces of rapamycin.

### Purification of the FRB0_EM_ construct

To generate a construct suitable for protein purification and cryo-EM structure determination, the FRB region of human mTOR (from residues 2021 to 2118) bearing the T2098L mutation was inserted into the TM2/TM3 loop of the *Danio rerio* TREK1 gene (drTREK1, K2P2.1, UniProt X1WC65). For biochemical stability, the drTREK1 gene was truncated to only include residues 1–322 of the full TREK1 ORF, and mutations to eliminate N-linked glycosylation sites (N95Q, N122Q) were inserted. The assembled FRB0_EM_ fusion construct was cloned into a pPICZ vector and fused to a PreScission protease-cleavage site (LEVLFQ/GP), followed by C-terminal GFP and His_10_ tags (Brohawn et al., 2014b) to facilitate purification. All genetic manipulation was performed by QuikChange mutagenesis or by SLIC cloning techniques. The pPICZ expression plasmid was linearized with a PmeI restriction enzyme and subsequently transformed by electroporation into an SMD1168H strain of *Pichia pastoris*. Screening for successful recombinant integration was performed by plating transformants on yeast extract peptone dextrose sorbitol (YPDS) plates containing increasing concentrations of zeocin from 0.5 mg/mL to 2 mg/ml.

Individual colonies that formed on zeocin plates were transferred into 10ml of buffered minimal medium (2 × YNB, 1% glycerol, 0.4 mg L−1 biotin, 100 mM potassium phosphate [pH 6.0]) in 50 mL conical tubes and incubated for 2 days at 30°C in a shaker at 220 rpm. The cells were pelleted by centrifugation (4000 × g at 20°C, 5 min) and resuspended in methanol minimal medium (2 × YNB, 0.5% methanol, 0.4 mg L−1 biotin, 100 mM potassium phosphate [pH 6.0]) to induce protein expression. Cells were shaken at 220 rpm for an additional 2 days at 22°C, with additional methanol added (final concentration 0.5% [v/v]) to the culture after 24 h of protein expression. After 48 h of protein expression, cells were pelleted by centrifugation (6500 × g at 4°C for 5 min), flash frozen in liquid nitrogen, and subjected to three rounds of cryo-milling (Retsch model MM301) in liquid N_2_ for 3 min at 25 Hz, to disrupt yeast cell walls and membranes. Frozen yeast cell powder was then stored at −80°C.

To purify FRB0_EM_ protein, cell powder was added to breaking buffer (150 mM KCl, 50 mM Tris pH 8.0, 1 mM phenylmethysulfonyl fluoride, 0.1 mg/mL DNase 1, and one tablet/50 mL of EDTA-free complete inhibitor cocktail [Roche]) buffer. Solubilized cell powder was centrifuged at 3000 × g at 4°C for 10 min to pellet large debris, and the supernatant was then centrifuged at 40,000 × g at 4°C for 1.5 h to pellet cell membranes. The pellet was re-suspended in 50 mL breaking buffer with 1 protease inhibitor tablet and 50 µL fresh PMSF 1 mM containing 60 mM n-Dodecyl-B-D-Maltoside (DDM) and incubated for 3 h with gentle stirring to solubilize the membranes, followed by centrifugation at 18500 × g at 4°C for 50 min. Talon cobalt resin (Takara Bio USA) was added to the supernatant at a ratio of 1 mL of resin per 10 g of cell powder and incubated in an orbital rotor overnight at 4°C. Resin was collected on a column and washed with 10 column volumes of low imidazole buffer (150 mM KCl, 50 mM Tris pH 8.0, 6 mM DDM, 30 mM imidazole) and bound protein was subsequently eluted from the resin by washing with high imidazole buffer (150 mM KCl, 50 mM Tris pH 8.0, 6 mM DDM, 300 mM imidazole). Homemade PreScission protease (∼1:25 wt:wt) was added to the eluate, and the cleavage reaction was allowed to proceed overnight at 4°C under gentle rocking. Cleaved FRB0_EM_ protein was concentrated in 50 kDa molecular weight cutoff (MWCO) Amicon Ultra Centrifugal Filters (Millipore) and applied to a Superdex200 10/300 gel filtration column (GE Healthcare) equilibrated in size exclusion chromatography (SEC) buffer (150 mM KCl, 20 mM Tris pH 8.0, 1 mM DDM). The purified FRB0_EM_ protein was concentrated (50 kDa MWCO) to 3 mg/mL and analyzed for purity by SDS-PAGE [12% (wt/vol) gels; Bio-Rad] followed by staining with Coomassie blue. All protein purification steps were carried out at 4°C.

### YFP/FKBP fusion protein purification and FRB0_EM_/FKBP complex formation

For bacterial expression of YFP/FKBP fusion protein, the human FKBP12 protein (Uniprot Q3V793) was cloned into a T7 promoter containing pET28 vector. A His_6_-YFP tag was inserted at the N-terminus of FKBP12 and a PreScission protease-cleavage site (LEVLFQ/GP) was inserted between the YFP and FKBP12 sequences. The YFP/FKBP pET28 plasmid was transformed into *Estrecheria coli* C41(DE3) cells (Lucigen), plated on LB plates supplemented with 50 µg/mL kanamycin, and grown overnight at 37°C. Following growth, all colonies were scraped from the plate using a rubber policeman and combined into a 25 mL pre-culture in kanamycin-supplemented LB media and grown overnight at 37°C in a shaker at 220 rpm. This pre-culture was then used to inoculate 4 liters of LB media supplemented with kanamycin and grow at 37°C in a shaker at 220 rpm. At an OD_600nm_ of 0.4, the temperature was reduced to 20°C and growth was continued until an OD_600nm_ of 0.8, at which point isopropyl-β-D-thiogalactoside (IPTG) was added to a final concentration of 0.5 mM to induce YFP-FKBP expression. Cells were then grown for an additional 12 h before being harvested by centrifugation (7,500g, 10 min, 4°C).

For purification of YFP/FKBP protein, all steps were performed at 4°C. Bacterial cell pellets were resuspended in 50 mL lysis buffer (150 mM KCl, 50 mM Tris, pH 8) supplemented with 1 mg lysozyme (Sigma-Aldrich), 1 mg DNase I (Sigma-Aldrich), 85 µg/mL PMSF (Roche), Leupeptin/Pepstatin (0.95/1.4 μg/mL, Roche), and 1 cOmplete ultra mini protease inhibitor tablet (Roche). Cells were lysed by sonication with a Sonic Dismembrator 500 (Thermo Fisher Scientific). Lysed cellular debris was cleared by centrifugation (18500 × g at 4°C for 50 min) and 5 mL of Talon cobalt resin (Takara Bio USA) was added to the supernatant and incubated in an orbital rotor overnight at 4°C. Resin was collected on a column and washed with 10 column volumes of low imidazole buffer (150 mM KCl, 50 mM Tris pH 8.0, 30 mM imidazole) and bound protein was subsequently eluted from the resin by washing with high imidazole buffer (150 mM KCl, 50 mM Tris pH 8.0, 300 mM imidazole). The eluted YFP/FKBP protein was concentrated in a 50 kDa molecular weight cutoff (MWCO) Amicon Ultra Centrifugal Filter (Millipore) and applied to a Superdex200 10/300 gel filtration column (GE Healthcare) equilibrated in size exclusion chromatography (SEC) buffer (150 mM KCl, 20 mM Tris pH 8.0). The peak fraction representing YFP-FKBP was collected and concentrated to 3 mg/mL.

A portion of the purified YFP/FKBP protein was flash frozen in liquid N_2_ and stored at −80 for fluorescent size exclusion chromatography studies to demonstrate protein complex formation between YFP/FKBP and the FRB0_EM_ protein in the presence of rapamycin (Figure 5a). The remaining protein was mixed with homemade PreScission protease (∼1:10 wt:wt) to cleave off the YFP and incubated for 3 h at 4°C, followed by a repeated run on a Superdex200 10/300 gel filtration column (GE Healthcare) equilibrated in size exclusion chromatography (SEC) buffer (150 mM KCl, 20 mM Tris pH 8.0). The purified FKBP protein was collected and concentrated to 3 mg/mL. To prepare the FRB0_EM_/rapamycin/FKBP ternary complex for cryo-EM studies, 250 µL of 4.9 mg/mL (51 µM) purified FRB0_EM_ protein was mixed with 250 µL of 2 mg/mL (46 µM) YFP-FKBP and 125 µL of 2 mg/mL 3C protease and incubated on ice for 2 h in the presence of 50 µM rapamycin. This mixture was then applied to a Superdex200 10/300 gel filtration column (GE Healthcare) equilibrated in size exclusion chromatography (SEC) buffer (150 mM KCl, 20 mM Tris pH 8.0). The peak fraction representing the FRB0_EM_/rapamycin/FKBP ternary complex was collected (Supplementary Figure 6b), concentrated to 3.1 mg/mL, and immediately frozen on cryo-EM grids.

### Fluorescent Size Exclusion Chromatography

Fluorescent size exclusion chromatography (FSEC) was used to assess for proper complex formation between the purified FRB and FKBP proteins in the presence of rapamycin. For samples analyzed by FSEC, 75 pmol of purified FRB0_EM_ was mixed with 35 pmol of purified YFP/FKBP fusion protein in 100 µL of size exclusion chromatography (SEC) buffer (150 mM KCl, 1 mM DDM, 20 mM Tris pH 8.0) containing either no rapamycin or an ascending range of rapamycin concentrations from 25 nM to 50 µM. Each sample was incubated for 30 min at 4°C to allow for complex formation and then applied to a Superdex 200 increase 5/150 GL gel filtration column (GE Healthcare) equilibrated in SEC buffer. Fluorescence signal was recorded using an in-line RF-20A XS fluorescence detector (Shimadzu) set for 514 nm excitation/ 529 nm emission to capture YFP fluorescence. Leftward shift of the YFP peak in rapamycin containing samples (from a retention volume of 2.71 mL to 2.12 mL) was indicative of complex formation. To estimate the *Ka* for complex formation, the ratio of the fluorescent signals at the two YFP peaks (2.12 mL / 2.71mL) was determined for each sample and normalized by the ratio value of the rapamycin free sample (Figure 5b).

### Cryo-EM grid preparation and data collection

Freshly purified FRB0_EM_ protein or FRB0_EM_/rapamycin/FKBP ternary complex protein samples was frozen on R1.2/1.3 UltraAUfoil 300 mesh grids (Quantifoil), prepared for sample application by glow discharge for 80 s at-25 mA using a Pelco easiGlow glow discharge system (Ted Pella). A 4 µL volume of 2.9 mg/mL purified TREK1 M2/M3 FRB0 protein or 3.1 mg/mL FRB0_EM_/rapamycin/FKBP complex was added to the grids and freezing was carried out using a Vitrobot Mark IV (FEI). The samples were equilibrated in the Vitrobot chamber at 21°C and 100% humidity for 20 s, blotted for 2 s with a blot force of +4, and then plunge frozen in liquid ethane.

The FRB0_EM_ datasets were acquired at 105,000x nominal magnification on a 300 kV Titan Krios G3i equipped with a Gatan K3 imaging system and a GIF quantum energy filter set to a slit width of 20 eV. Data were collected using Leginon v 3.6 software^72^, at a dose rate of 42.06 e^-^/Å^2^/s with a total exposure of 1.40 s, for an accumulated dose of 58.88 e^-^/Å^2^. Intermediate frames were collected every 0.04 s, for a total of 35 frames per micrograph. A total of 5967 images were collected for the FRB0_EM_ sample frozen in the absence of rapamycin, at a nominal defocus range of 0.6 – 2.5 µm. An additional 3147 images were then collected from the FRB0_EM_ sample in the presence of rapamycin, at a nominal defocus range of 0.7 – 2.8 µm. For both datasets, a calibrated pixel size of 0.4222 Å/pixel was used for data processing.

The FRB0_EM_/rapamycin/FKBP ternary complex dataset was acquired at 105,000x nominal magnification on a 300 kV Titan Krios G3i equipped with a Gatan K3 imaging system and a GIF quantum energy filter set to a slit width of 20 eV. Data were collected using Leginon v 3.7 software^72^, at a dose rate of 24.94 e^-^/Å^2^/s with a total exposure of 1.80 seconds, for an accumulated dose of 44.90 e^-^/Å^2^. Intermediate frames were collected every 0.05 s, for a total of 40 frames per micrograph. A total of 12,397 images were collected at a nominal defocus range of 0.6 – 2.7 µm, with a calibrated pixel size of 0.4125 Å/pixel used for data processing.

### CryoEM data processing

For processing of the FRB0_EM_ data set, movie frames from 5,967 micrographs were aligned using Patch Motion Correction in cryoSPARC v.4.5.3^73^ with a B-factor during alignment of 500. Contrast transfer function (CTF) estimation was performed with Patch CTF in cryoSPARC v.4.5.3. Particles were picked using blob picker in cryoSPARC, followed by several rounds of 2D classification to generate 2D classes, which were then used as input for template picker. The initial template picker yielded 5,172,684 particles, which were extracted in cryoSPARC with a 496-pixel box size, binned four times, and then classified via 2D classification in cryoSPARC v.4.5.3. Well-defined, high-resolution 2D classes were selected, resulting in a particle stack of 1,038,638 particles. These particles underwent further sorting through 2D classification with a “Batchsize per class” of 400, “Force Max over poses/shifts” turned off, and 40 online-EM iterations. This process reduced the particle stack to 88,719 particles, which were then subjected to a three-class ab initio reconstruction with a maximum resolution of 7 Å and an initial resolution of 9 Å. One class displayed features consistent with the TREK1 protein, while the other two lacked them (Supplementary Fig. 7). We returned to the particle stack of 1,038,638 particles and performed heterogeneous refinement in cryoSPARC, using the volume from the best class in the ab initio reconstruction, an alternative ab initio volume as a decoy, and three additional decoy classes. We set the “Batch size per class” to 30,000 and the “Number of final full iterations” to 5. The particles from the best class of this heterogeneous refinement (217,166 particles) were next fed into a three-class ab initio refinement (as above) to improve the input volume for subsequent heterogeneous refinements. We repeated this process for three rounds, alternating heterogeneous refinements with decoy classes and ab-initio refinements, in each instance using a smaller subset of particles from the good class obtained by heterogenous refinement as input for the next round. The good input volume for each round of heterogeneous refinement was updated after each ab initio reconstruction, with a non-uniform refinement job used to achieve the highest possible resolution reconstruction prior to advancing to the next round. After three rounds, the particle stack for the good volume contained 38,923 particles (Supplementary Fig. 7). These particles were then re-extracted without binning in a box size of 496 pixels. The un-binned particles were used as input for a single-class ab initio reconstruction, with a maximum resolution set to 3 Å and an initial resolution of 5 Å, using “Fourier radius step” of 0.005, a “Final minibatch size” of 1000, and the “Center structures in real space” turned off^74^. The resulting ab initio particle stack was used for 3D reconstruction with “Force re-do GS split” enabled. Finally, the particles, volume, and mask from the 3D reconstruction were used as input for local refinement, with a rotation search extent of 2 degrees, a shift search extent of 1, “Re-center rotations” turned on each iteration, and “Re-center shifts” turned on each iteration, achieving a final resolution of 4.33 Å.

For processing of the FRB0_EM_/rapamycin/FKBP ternary complex data set, movie frames from 12,398 micrographs were aligned using Patch Motion Correction in cryoSPARC v.4.2.1^73^ with a B-factor during alignment of 500. Contrast transfer function (CTF) estimation was performed with Patch CTF in cryoSPARC v.4.2.1. Particles were picked using blob picker in cryoSPARC, followed by several rounds of 2D classification to generate 2D classes, which were then used as input for template picker in cryoSPARC v4.2.1. The initial template picker yielded 2,338,297 particles, which were extracted in cryoSPARC with a 480-pixel box size, without binning. These particles were sorted using 2D classification in cryoSPARC, resulting in a particle stack of 346,637 particles which were used as input for a three-class ab initio reconstruction with a maximum resolution set to 3 Å and an initial resolution of 5 Å, using “Fourier radius step” of 0.005, a “Final minibatch size” of 1000, and the “Center structures in real space” turned off^74^ (Supplementary Fig. 8). The class displaying features of both TREK and density outside the membrane was selected, and the three-class ab initio reconstruction was repeated, resulting in a particle stack of 53,477 particles. This particle stack was subjected to 3D classification with 10 classes using a mask designed to exclude the detergent micelle and include all protein density, with a target resolution of 7 Å. The “O-EM batch size (per class)” was set to 3000, the “O-EM learning rate init” to 1, and the “O-EM learning rate halflife (%)” to 0. The classes were sorted into two groups based on similarity; the particle stack from one group was used as input for a single-class ab initio reconstruction with a maximum resolution of 3 Å and an initial resolution of 5 Å, using a “Fourier radius step” of 0.005, a “Final minibatch size” of 1000, and the “Center structures in real space” turned off (Supplementary Fig. 8). The resulting ab initio particle stack was used for 3D reconstruction with “Force re-do GS split” enabled. Finally, the particles, volume, and mask from the 3D reconstruction were used as input for local refinement, with a rotation search extent of 2 degrees, a shift search extent of 1, and “Re-center rotations” and “ Re-center shifts” turned on each iteration, achieving a final resolution of 4.20 Å.

We also achieved a higher resolution (3.60 Å) map from a subset of particles in this dataset where the density for FRB and FKBP was not visible. For this, we focused on the class that displayed density for TREK1 but not for the FRB/FKBP in the initial three-class ab initio reconstruction (Supplementary Figure 8c). Particles from this “TREK1-only” class were used as input for a 3D reconstruction with “Force re-do GS split” enabled, and finally, the particles, volume, and mask from the 3D reconstruction were used as input for local refinement, with a rotation search extent of 2 degrees, a shift search extent of 1, “Re-center rotations” turned on each iteration, and “Re-center shifts” turned on each iteration, achieving a final resolution of 3.63 Å. This reconstruction appeared to contain a heterogeneous mixture of particles. The TM4 helix in one subunit was consistently positioned in the “up” conformation, while the TM4 helix in the second subunit was positioned in a mixture of “up” or “down” conformations. To sort out this heterogeneity, we ran an additional three-class ab initio reconstruction, followed by 3D reconstruction and local refinement of the two best classes, resulting in two maps at 3.74 Å (TM4 up) and 3.64 Å (TM4 down).

---

**Supplemental Figure 1:**
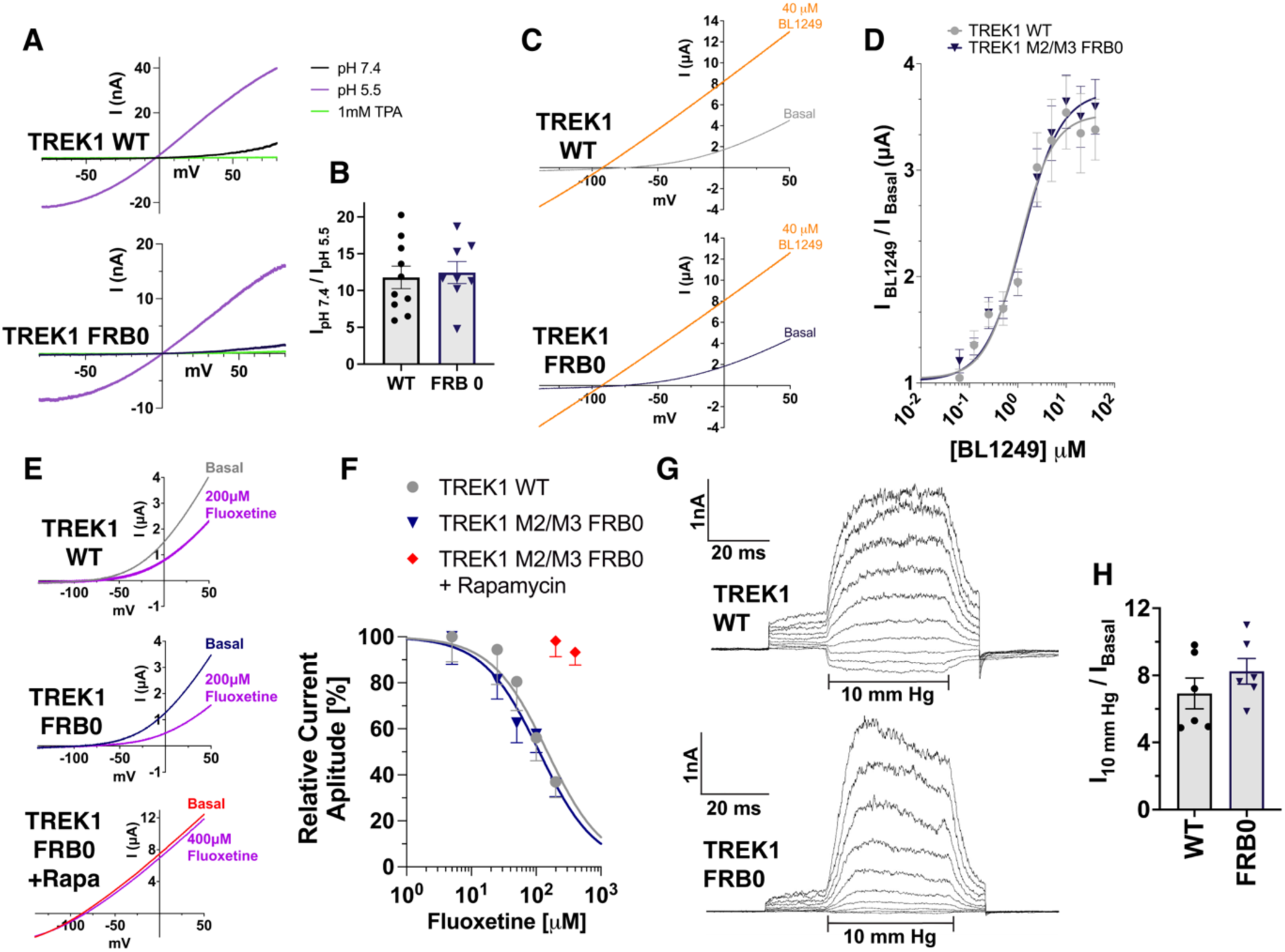
Gating behavior of the TREK1 FRB fusion channel. **A.** Exemplar *X. laevis* oocyte inside-out patch clamp recordings of TREK1 WT (upper panel) or TREK1 FRB0 (lower panel) channels, first perfused with buffer at pH 7.4 (black trace), then switched to pH 5.5 (blue trace), and then to pH 5.5 in the presence of 1mM tetrapentylammonium (TPA, yellow trace). **B.** Quantitation of normalized current responses to treatment with pH 5.5, for TREK1 WT or TREK1 FRB channels. **C.** Exemplar *X. laevis* oocyte TEVC recordings of TREK1 WT (upper panel) or TREK1 FRB0 (lower panel) channels to treatment with 40 µM BL1249 and **D.** dose response curves showing normalized response to treatment with BL1249. **E.** Exemplar *X. laevis* oocyte TEVC recordings of TREK1 WT (upper panel), TREK1 FRB0 (middle panel), or TREK1 FRB0 after treatment with 2.5 µM rapamycin (lower panel) to treatment with fluoxetine and **F.** dose response curves showing normalized response to treatment with fluoxetine. **G.** Exemplar *X. laevis* oocyte inside-out patch clamp recordings of TREK1 WT (upper panel) or TREK1 FRB0 (lower panel) channels, exposed to voltage steps from −80 to +100 mV in 20 mV increments, prior to and after application of −10 mm Hg pressure to the patch pipette and **H.** Quantitation of normalized current responses to treatment with −10 mm Hg pressure, for TREK1 WT or TREK1 FRB channels.

**Supplemental Figure 2:**
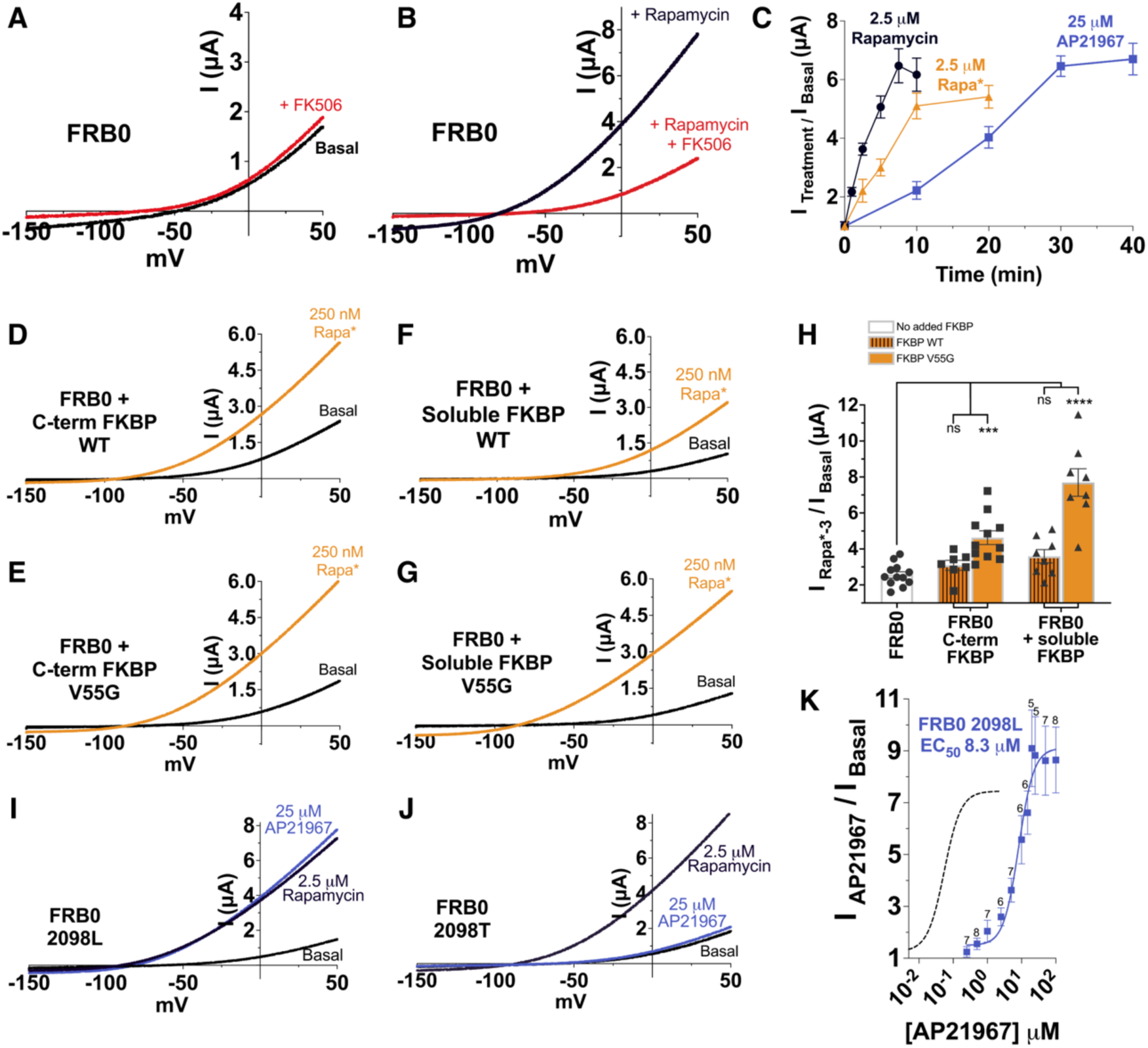
Modulation of the TREK1 FRB0 channel by Rapamycin analogs. A,B. Exemplar *X. laevis* oocyte TEVC recordings of TREK1 FRB0 channels, after treatment with either 2.5 µM FK506 (**A**, red trace), 2.5 µM rapamycin (**B**, blue trace), or a 10-minute pretreatment with 2.5 µM FK506 followed by co-incubation with 2.5 µM concentrations of both rapamycin and FK506 (**B**, red trace). **C.** Time course of FRB0 channel activation by rapamycin, Rapa*, and AP21967. For all time points, current at 0 mV after drug treatment was normalized against paired untreated oocytes. **D-G.** Exemplar *X. laevis* oocyte TEVC recordings, showing response to a 20-minute treatment with 250 nM Rapa*, for (**D**) TREK1 FRB0 C-terminal FKBP, (**E**) TREK1 FRB0 C-terminal FKBP V55G, or TREK1 FRB0 co-expressed with soluble cytoplasmic (**F**) FKBP WT or (**G**) FKBP V55G, **H.** Quantitation of normalized current responses to treatment with 250 nM Rapa*, for the FRB0 channel alone (white bar) or for FRB0 channels in the presence of FKBP WT (hatched orange bars) or FKBP V55G mutant (solid orange bars). **I,J.** Exemplar *X. laevis* oocyte TEVC recordings of (**I**) TREK1 FRB0 2098L channels or (**J**) TREK1 FRB0 2098T channels, after a 10-minute incubation with 2.5 µM rapamycin or a 40-minute incubation with 25 µM AP21967. **K.** AP21967 dose response relationship for the TREK1 FRB0 channel. Number of replicates for each point is shown; error bars represent SEM. For comparison, dashed curve reproduces the TREK1 FRB0 channel dose response relationship after rapamycin treatment, as shown in Figure 1E.

**Supplemental Figure 3:**
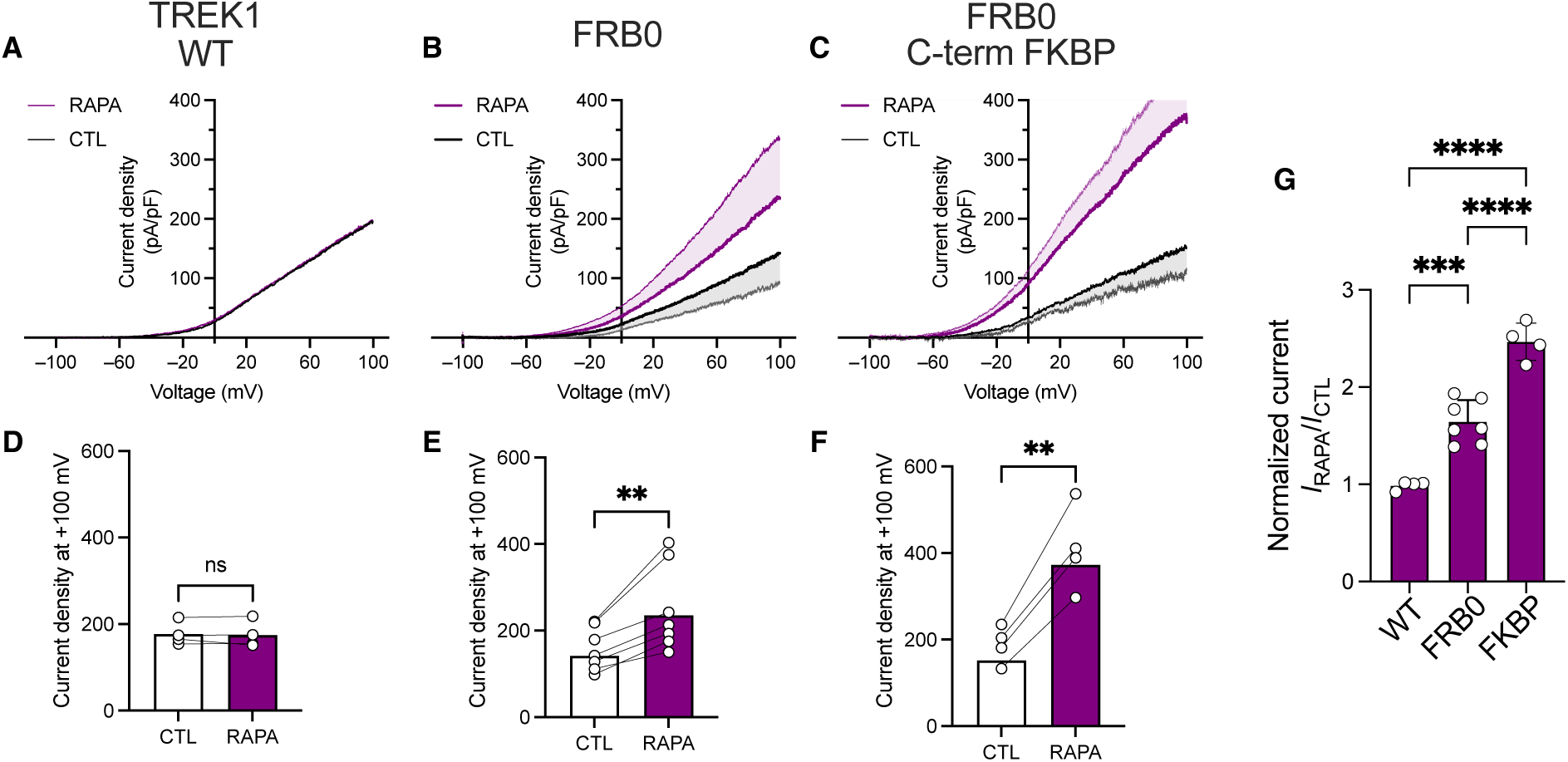
Modulation of TREK1 FRB fusion channels in a mammalian cell background. Exemplar traces from whole cell patch clamp recordings of TREK1 WT **(A)**, TREK1 FRB0 **(B)**, or TREK1 FRB0 C-terminal FKBP channels **(C)**, before (CTL) or following treatment with 2.5 µM rapamycin (RAPA). **D,E,F.** Quantitation of current density at +100mV, before or following application of 2.5 µM rapamycin to **(D)** TREK1 WT, **(E)** FRB0, or **(F)** TREK1 FRB0 C-terminal FKBP channels **G.** Quantitation of normalized current density prior to and following rapamycin treatment, for TREK1 WT (WT), TREK1 FRB0 (FRB0), and TREK1 FRB0 C-terminal FKBP (FKBP). Number of replicates for each point is shown. Statistical significance was determined by one-way ANOVA, combined with a Tukey multiple post-hoc comparison against TREK1 WT data. Results indicated, **p<0.05, ***p<0.005, ****p<0.0005. Error bars are mean ± SD.

**Supplemental Figure 4:**
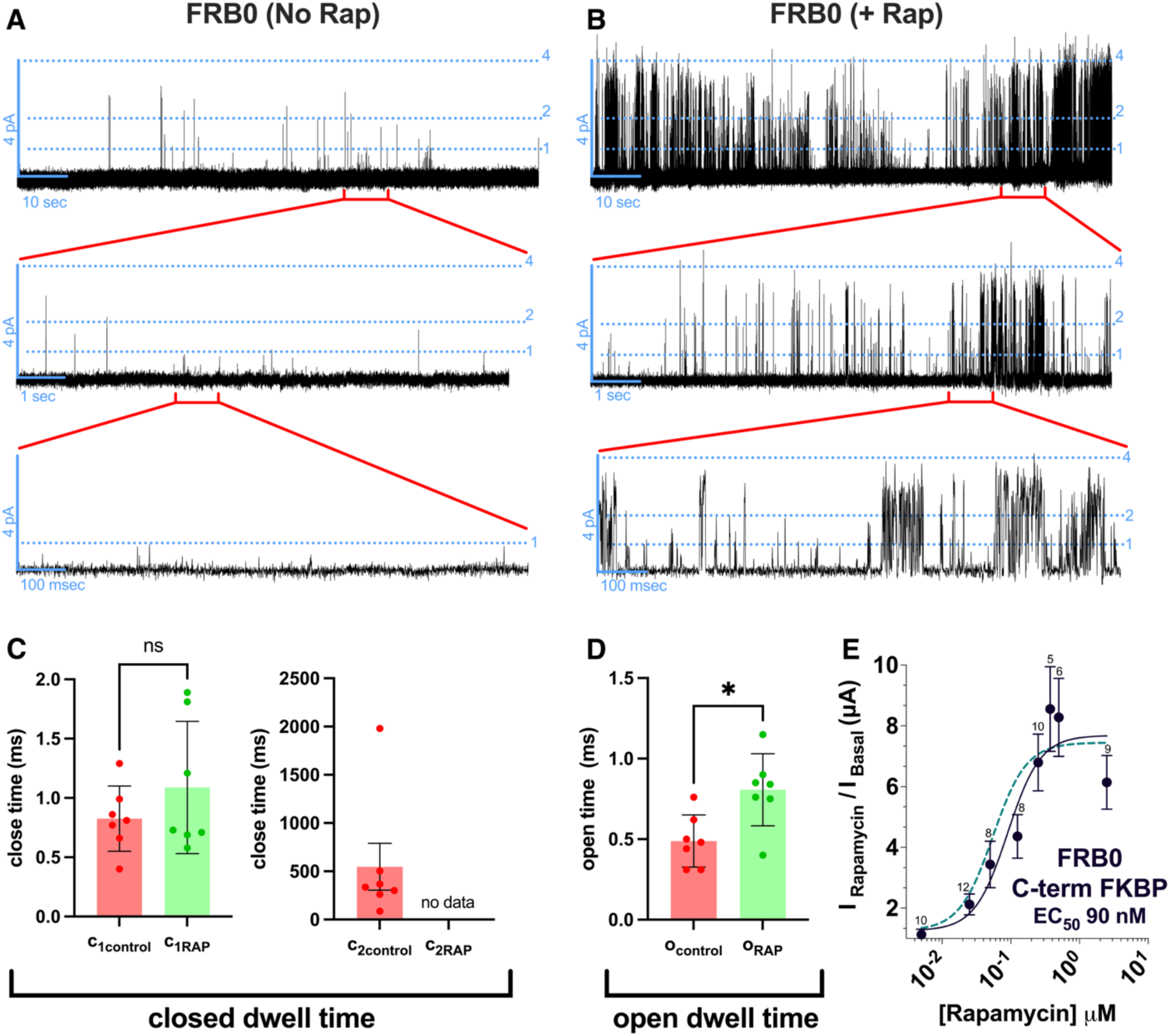
Single channel behavior of the FRB0 channel A,B. Representative recording of a single TREK1 FRB0 C-term FKBP channel (A) prior to and (B) following addition of 25 µM rapamycin. Upper panels show a continuous 100-s record. Middle panels show a select 10-s region of the upper panel, at the region noted. Lower panels show a select 1-s region of the middle panel, at the region noted. Lower panels are reproductions of those shown in Figure 3A. **C,D.** Quantitation of mean (C) closed and (D) open dwell times for each individual single-channel recording. Number of replicates for each point is shown. Statistical significance was determined by Wilcoxson signed rank test. Results indicated, *p<0.03, Error bars are mean ± SD. **E.** Rapamycin dose response relationship for the TREK1 FRB0 C-term FKBP channel, measured by two electrode voltage clamp experiments. Number of replicates for each point is shown; error bars represent SEM. For comparison, dashed grey curve reproduces the TREK1 FRB0 channel dose response relationship after rapamycin treatment, as shown in Figure 1E.

**Supplemental Figure 5:**
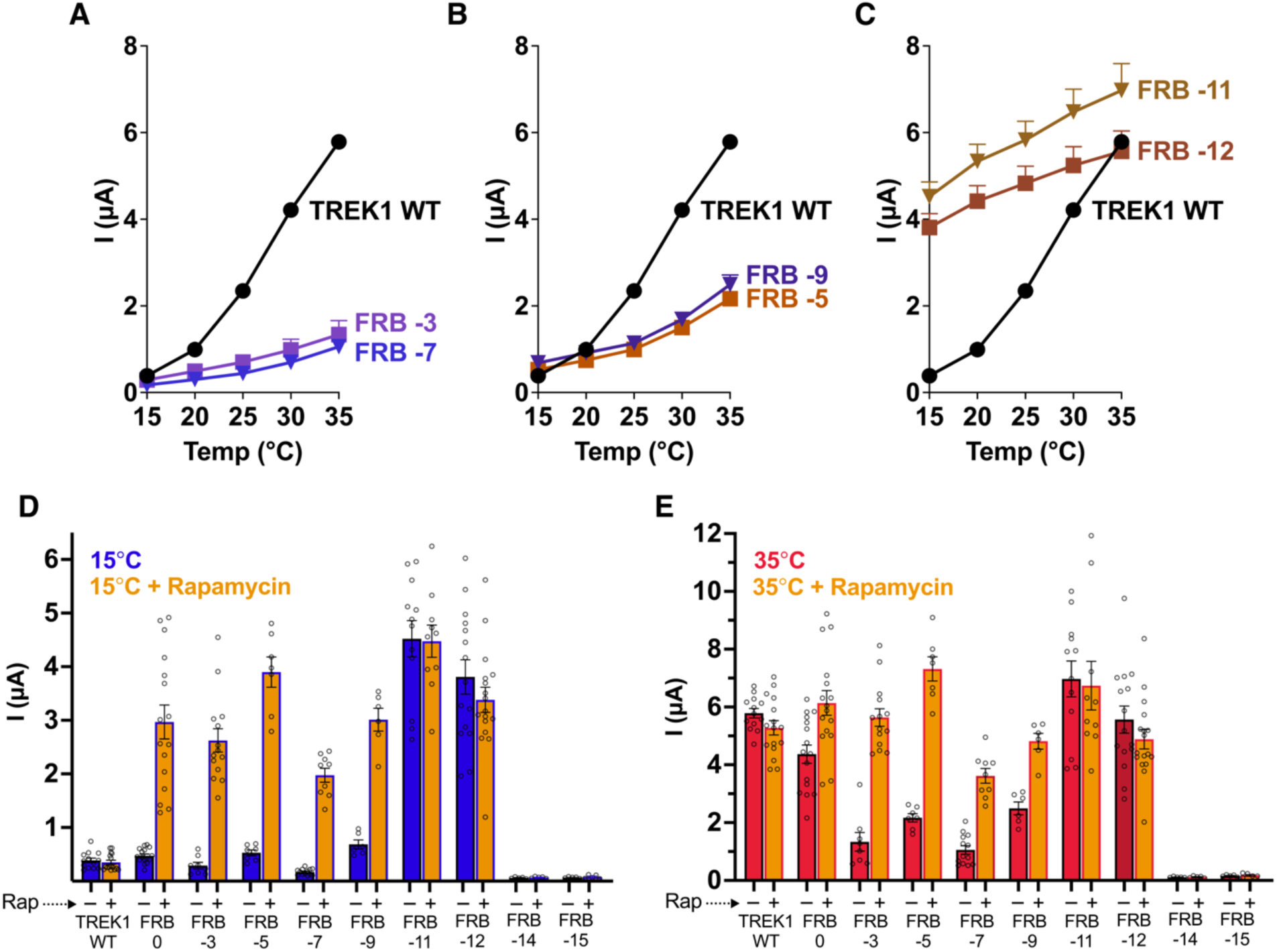
Temperature responsiveness of TREK1 FRB0 channels with alternative TM2 fusion sites. A-C. Quantification of TREK1 wildtype or FRB-n construct responses to temperature, as measured by TREK1 current at 0 mV, for (**A**) FRB-n mutants that exhibit diminished activation by heat, (**B**) FRB-n mutants that exhibit both diminished inhibition by cold and activation by heat, or (**C**) FRB-n mutants that exhibit diminished inhibition by cold. **D-E.** TREK1 wildtype or FRB-n construct currents at 0 mV in either the absence or presence of 2.5 µM rapamycin, measured at (**D**) 15°C or (**E**) 35°C.

**Supplemental Figure 6:**
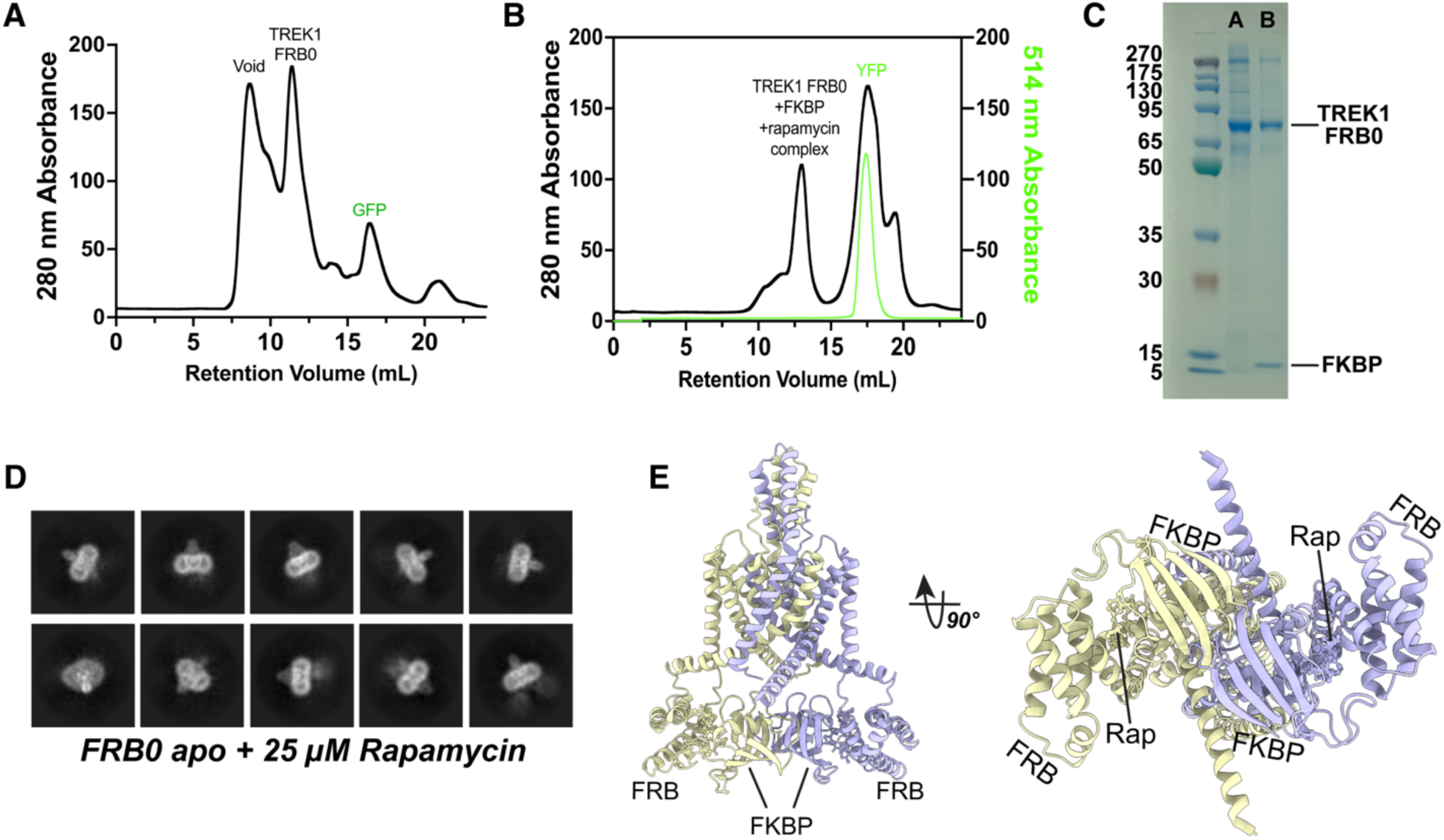
Biochemical properties of the FRB0 channel and FRB0/rapamycin/FKBP complex. **A.** Size exclusion chromatography profile of the TREK1 FRB0 channel after 3C cleavage of the GFP purification tag. **B.** Size exclusion chromatography profile of the rapamycin-induced complex between the TREK1 FRB0 channel and FKBP. Absorbance at 280 nm (black, left axis) and 514 nm (yellow, right axis) demonstrate complete removal of the YFP tag from the FKBP in the FRB0/FKBP complex. **C.** SDS-PAGE of the TREK1 FRB0 samples collected as indicated in (A) and the TREK1 FRB0/FKBP/rapamycin complex collected as indicated in (B). **D.** Representative 2D classes of TREK1 FRB0 channel protein incubated with 25 µM rapamycin prior to freezing. **E.** A hypothetical model of the TREK1 FRB0 channel in-complex with two FKBP subunits, created by duplicating the model of the FKBP-bound subunit (as shown in Figure 5G) and aligning this duplicated model with the position of the free subunit in the experimental data. In the bottom view panel (right), a clash between the two bound FKBP molecules is apparent.

**Supplemental Figure 7:**
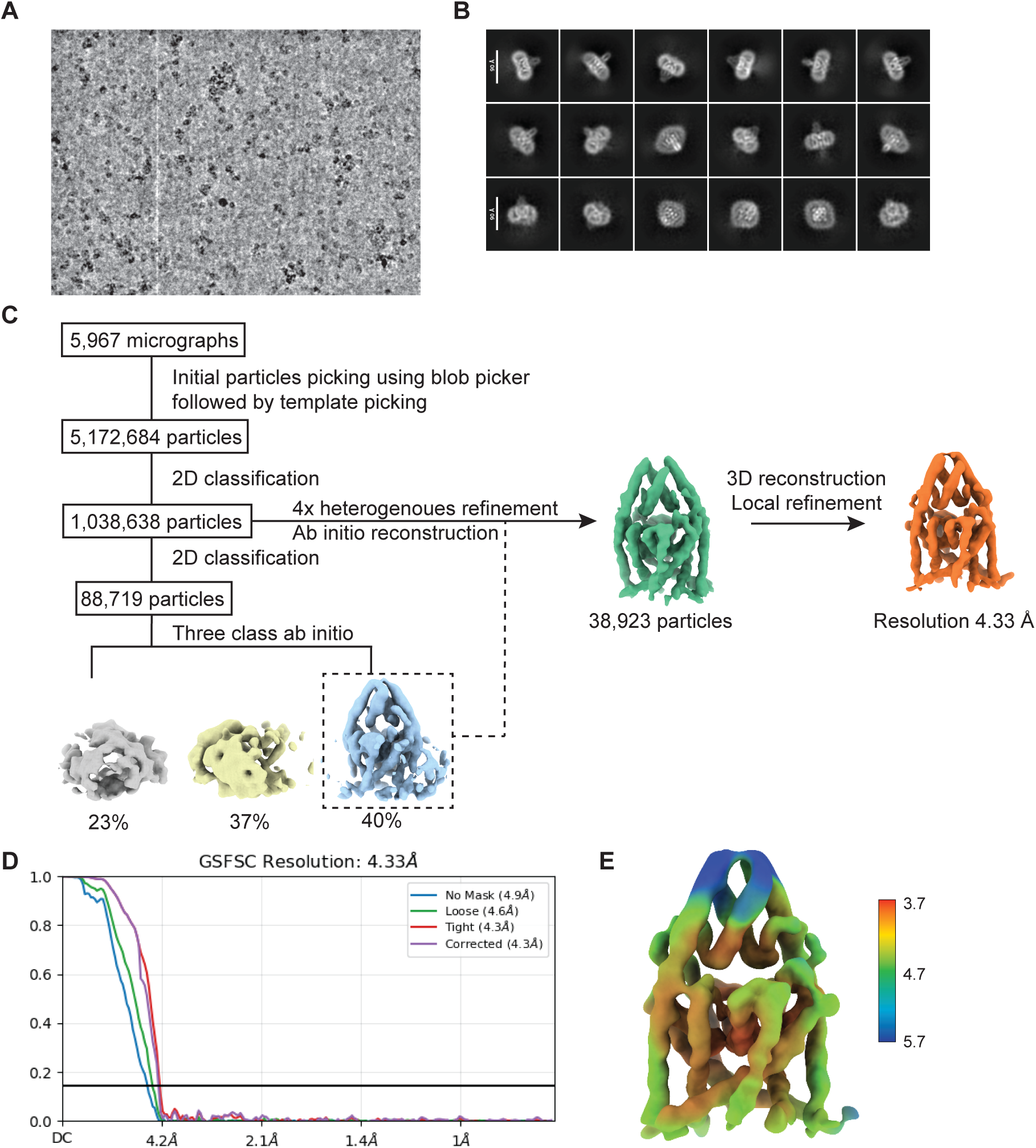
Cryo-EM processing workflow for the FRB0 channel in the absence of rapamycin or FKBP. Representative **A.** micrograph from the dataset and **B.** 2D classes of the FRB0 channel. **C.** Overview of the processing scheme, highlighting major branch points in the processing pipeline**. D.** cryoSPARC generated FSC curve from the final refinement of the FRB0 map. **E.** cryoSPARC generated local resolution map.

**Supplemental Figure 8:**
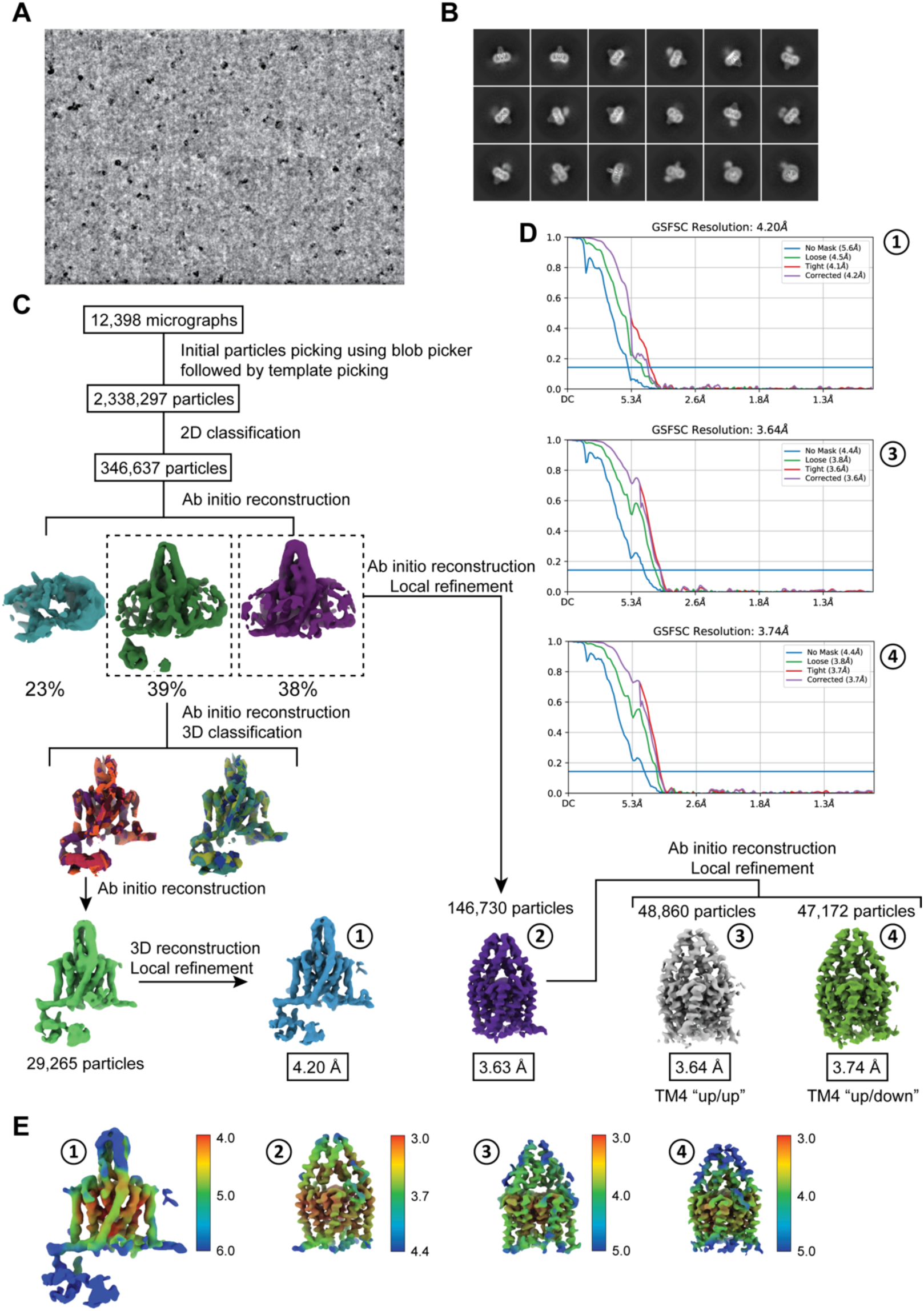
Cryo-EM processing workflow for the FRB0 channel in complex with rapamycin and FKBP. Representative **A.** micrograph from the dataset and **B.** 2D classes of the FRB0/rapamycin/FKBP complex. **C.** Overview of the processing scheme, highlighting major branch points in the processing pipeline**. D.** cryoSPARC generated FSC curve from the final models generated in this dataset. **E.** CryoSPARC generated local resolution maps.

**Supplemental Figure 9:**
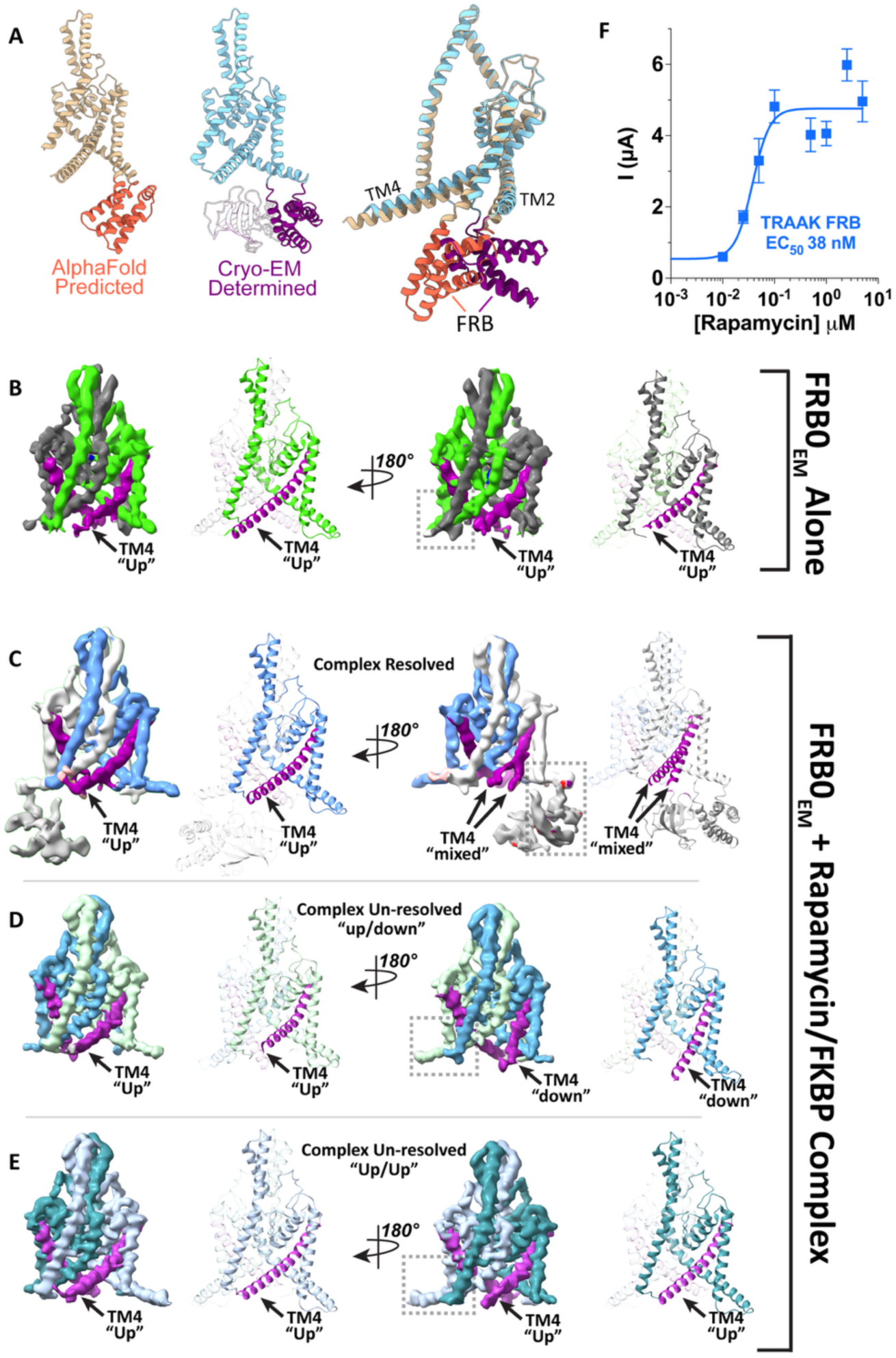
TM4 positioning in the FRB0 channel cryo-EM maps. **A.** Comparison of an AlphaFold model of the FRB0 channel (left) and the experimentally determined map of FRB0 in complex with rapamycin and FKBP (middle) and an overlay of the two models (right). In the center panel, the rapamycin and FKBP are shown but transparent, to highlight comparison of the position of the FRB domain. **B.** Final unsharpened cryo-EM map of the FRB0 channel, with the TM4 helix highlighted in purple. **C, D, E** Equivalent representations of the final unsharpened cryo-EM maps of the FRB0 channel in the presence of rapamycin and FKBP when **C.** the complex is resolved in the map, **D.** the complex is un-resolved and the channel is in a TM4 “up/down” conformation, or **E.** the complex is un-resolved and the channel is in a TM4 “up/up” conformation. **F.** Rapamycin dose response relationship for the TRAAK FRB channel, measured by two electrode voltage clamp experiments. Number of replicates for each point is shown; error bars represent SEM.

**Supplemental Figure 10:**
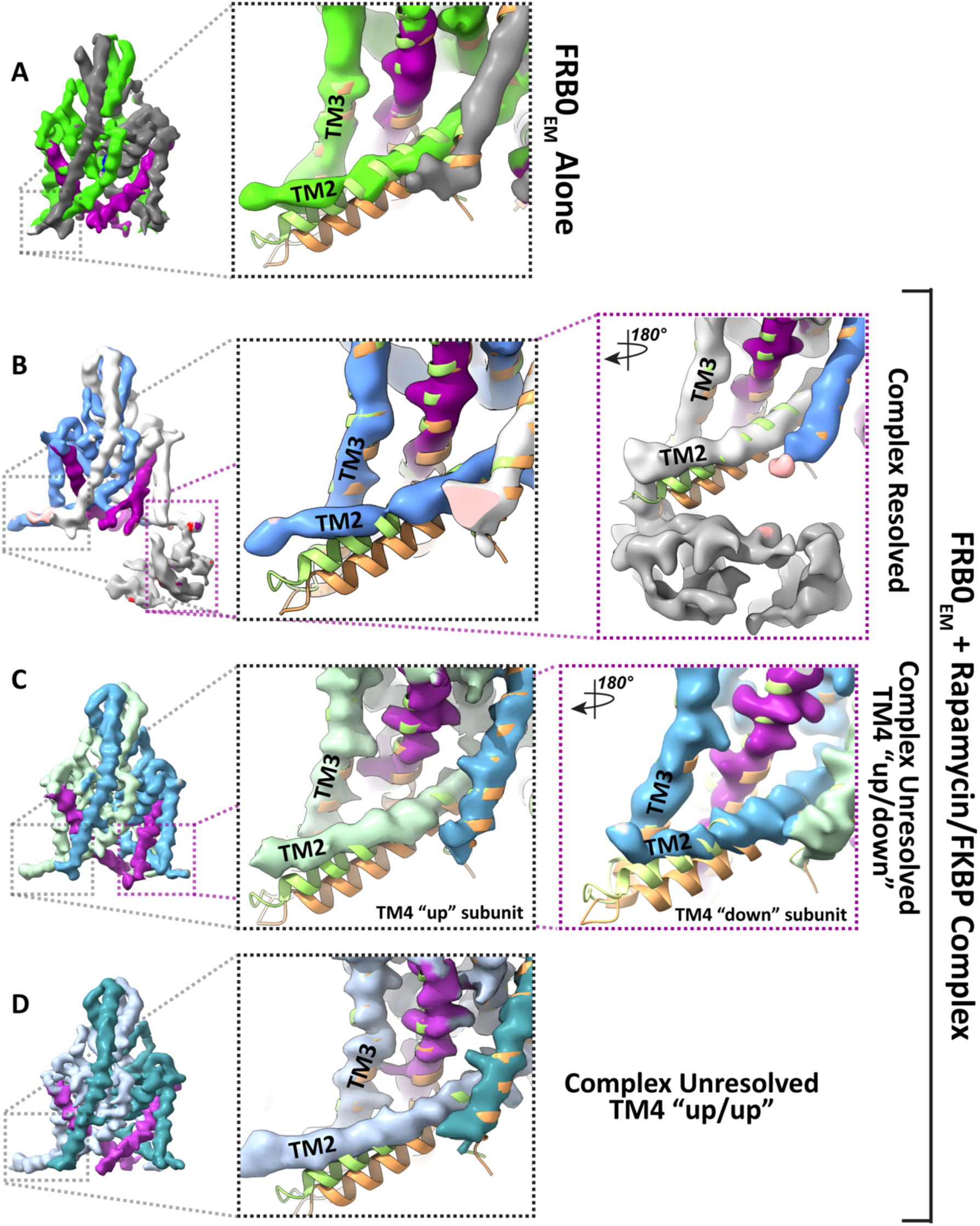
FRB0 insertion alters the position of the TM2 helix. Final unsharpened cryo-EM maps of **A.** the FRB0 channel in the absence of rapamycin and FKBP, **B.** the FRB0 channel in the presence of rapamycin and FKBP complex, or **C,D.** the maps from the sample of the FRB0 channel in the presence of rapamycin and FKBP complex where the complex is unresolved. Enlarged insets (right) show the TM2/TM3 loop region overlayed on previously determined models of TREK1^38^ in the TM4 “up” (yellow model, PDB: 8DE8) or TM4 “down” conformations (orange model, PDB: 8DE9).

## Notes

### Competing Interest Statement

The authors have declared no competing interest.

### Funding

- NIH Common Fund: R01GM145918

